# Exploring the impact of read clustering thresholds on RADseq-based systematics: an empirical example from European amphibians

**DOI:** 10.1101/2023.04.19.537466

**Authors:** Loïs Rancilhac, Florent Sylvestre, Carl R. Hutter, Jan W. Arntzen, Wieslaw Babik, Pierre-Andre Crochet, Grégory Deso, Rémi Duguet, Pedro Galan, Maciej Pabijan, Mathieu Policain, Pauline Priol, Joana Sabino-Pinto, Maria Capstick, Kathryn R. Elmer, Christophe Dufresnes, Miguel Vences

## Abstract

Restriction site-Associated DNA sequencing (RADseq) has great potential for genome-wide systematics studies of non-model organisms. However, accurately assembling RADseq reads into orthologous loci remains a major challenge in the absence of a reference genome. Traditional assembly pipelines cluster putative orthologous sequences based on a user-defined clustering threshold. Because improper clustering of orthologs is expected to affect results in downstream analyses, it is crucial to design pipelines for empirically optimizing the clustering threshold. While this issue has been largely discussed from a population genomics perspective, it remains understudied in the context of phylogenomics and coalescent species delimitation. To address this issue, we generated RADseq assemblies of representatives of the amphibian genera *Discoglossus, Rana, Lissotriton* and *Triturus* using a wide range of clustering thresholds. Particularly, we studied the effects of the intra-sample Clustering Threshold (iCT) and between-sample Clustering Threshold (bCT) separately, as both are expected to differ in multi-species data sets. The obtained assemblies were used for downstream inference of concatenation-based phylogenies, and multi-species coalescent species trees and species delimitation. The results were evaluated in the light of a reference genome-wide phylogeny calculated from newly generated Hybrid-Enrichment markers, as well as extensive background knowledge on the species’ systematics. Overall, our analyses show that the inferred topologies and their resolution are resilient to changes of the iCT and bCT, regardless of the analytical method employed. Except for some extreme clustering thresholds, all assemblies yielded identical, well-supported inter-species relationships that were mostly congruent with those inferred from the reference Hybrid-Enrichment data set. Similarly, coalescent species delimitation was consistent among similarity threshold values. However, we identified a strong effect of the bCT on the branch lengths of concatenation and species trees, with higher bCTs yielding trees with shorter branches, which might be a pitfall for downstream inferences of evolutionary rates. Our results suggest that the choice of assembly parameters for RADseq data in the context of shallow phylogenomics might be less challenging than previously thought. Finally, we propose a pipeline for empirical optimization of the iCT and bCT, implemented in optiRADCT, a series of scripts readily usable for future RADseq studies.

## 1. Introduction

In the past three decades, the tools available for molecular phylogenetics, phylogeography and systematics have been rapidly evolving. Recently, High-Throughput Sequencing (HTS) methods, coupled with genome reduced-representation approaches, have enabled sequencing numerous genetic markers in many samples at reasonable cost, facilitating the transition from single-locus via multi-locus to genome-wide studies (Lemmon & Lemmon 2013; McCormack et al. 2013). Consequently, inference of evolutionary history has gained accuracy from the improved informativeness of the data and a more comprehensive representation of the heterogeneity of the phylogenetic signal across loci. However, HTS-based approaches also bring new challenges at the steps of data generation, assembly, filtering, and in downstream analyses (Scornavacca et al. 2020). Especially, issues related to data assembly and filtering have received widespread attention as potential sources of bias in downstream analyses, a problem that may be exacerbated when studying non-model organisms without reference genomes, and/or anonymous markers. In this context, it is important to be able to predict the impact of bias introduced by improper assembly of HTS data and to use specific approaches to optimize assembly steps.

Restriction site-Associated DNA sequencing (RADseq; Miller et al. 2007) has become one of the most widely used genome reduced-representation approaches (Andrews et al. 2016), with a large variety of derived protocols (e.g., genotyping-by-sequencing, Elshire et al. 2011; double-digest RADseq, Peterson et al. 2012). Although initially developed for population-level studies, the relevance of data obtained via RADseq and related approaches (hereafter subsumed under the term RADseq) for phylogenetic inference was emphasized early on (Cariou et al. 2013), even at relatively deep evolutionary scales (Herrera & Shank 2016). The success of this approach can be attributed to its relatively low cost, large number of unlinked markers generated, and applicability to non-model organisms. RADseq data can be assembled into loci by mapping to a reference genome when available or, more commonly, assembled *de novo*. *De novo* assembly of RADseq data usually follows three main steps (Eaton 2014): 1) the reads are grouped into clusters within every sample independently, and a consensus sequence is generated for each cluster; 2) the consensus sequences are assembled into presumably orthologous loci across individuals; 3) outlier loci are filtered out and the data is formatted for downstream analyses. For the first two steps, the user must set a similarity threshold (hereafter referred to as Clustering Threshold [CT] as in ipyrad, Eaton & Overcast 2020; corresponding to parameters *M* and *n* in stacks, Rochette et al. 2019) above which the reads and consensus sequences, respectively, are considered homologous. This parameter is particularly important because misspecifications might strongly impact downstream analyses (McCartney-Melstad et al. 2019). If the value is too low, non-orthologous sequences might be clustered together (“over-merging” following the terminology of Hühn et al. 2022; sometimes referred to as “over-lumping”), which would artificially increase the heterozygosity of the assembled loci. Filters can be set to entirely remove over-variable clusters and loci, but this will result in a loss of data that might otherwise have been clustered properly. Conversely, a CT too conservative might split alleles of polymorphic loci into independent loci (“under-merging”, also sometimes referred to as “over-splitting”), resulting in an underestimation of genetic diversity, and overall biases toward slowly evolving loci in final assemblies. In addition, more clusters of lower depth will be built, increasing the number of clusters lost to depth filters, and the computational burden of the assembly. Thus, one major challenge when assembling RADseq data is to identify a correct CT that maximizes the informativeness of the assembly while avoiding the clustering of non-orthologous sequences.

Several studies have explored strategies to determine the best fitting CT for a given data set, usually by assembling the same raw data with a range of CTs and comparing various characteristics of the different assemblies (reviewed in McCartney-Melstad et al. 2019; see also Ilut et al. 2014; Mastretta-Yanes et al. 2015; Paris et al. 2017). However, the applicability of such approaches to multi-species data sets assembled for phylogenetic studies remains limited for two main reasons. First, they often rely on the sequencing of many conspecific individuals (e.g., technical replicates or multiple individuals from a single locality in Mastretta-Yanes et al. 2015; isolation by distance slopes in McCartney-Melstad et al. 2019), whereas phylogenetic data sets often include only a single or few individuals per species. Secondly, an often-overlooked difference between single- vs. multi-species studies is that the former generally use the same CT to cluster reads within samples and consensus sequences among samples (Paris et al. 2017; Gagnaire 2020). While this might be appropriate when considering closely related conspecific individuals, as one would not expect alleles from a single individual to be substantially more divergent than two alleles randomly drawn from the population, such an assumption could result in incorrect clustering when considering species at higher divergence levels. Thus, assembly of multi-species RADseq data should rely on adapted protocols to optimize the CT within- and between-sample separately (Karbstein et al. 2020; Hühn et al. 2022).

Given the multiple effects of inadequate CT specification, both concatenation-based and single-gene inferences of phylogeny might be greatly impacted. If the CT is not conservative enough, paralogs could be clustered together, resulting in erroneous phylogenetic signal in both concatenation trees (Fitch 2000) and gene genealogies (Degnan & Rosenberg 2009). An overly conservative CT would result in splitting genuine heterozygous loci, reducing the amount of phylogenetic information within the data. Multi-Species Coalescent (MSC) inferences would also be affected by artefacts in gene genealogies. In both cases, the estimation of the topology and the branch lengths of the phylogenetic tree could be impacted, but the evidence for this effect is mixed. Rubin et al. (2012) found in an *in silico* analysis that the informativeness of the assemblies was a stronger driver of accuracy in RADseq based phylogenetic inference than an optimal orthology assessment. Accordingly, empirical studies in various taxa recovered topologies that were resilient to changes in the CT (e.g., Cruaud et al. 2014; Rancilhac et al. 2019; Piwczyński et al. 2021). On the other hand, Leaché et al. (2015a) reported important conflicts among phylogenetic trees reconstructed with various CTs, using both concatenation and species tree approaches. Recently, assembly metrics have been used for optimizing both within- and between-sample CTs in the context of phylogenetic studies (e.g., Paetzold et al. 2019; Karbstein et al. 2020; Hühn et al. 2022). One potential pitfall of this approach is that assembly metrics do not provide a direct assessment of orthology, meaning that perceived improvements do not necessarily correspond to a better clustering of orthologs. Especially, in the absence of *a priori* knowledge of the phylogenetic relationships of the studied taxa, it is difficult to assess whether improved support in downstream phylogenetic analyses following metrics-based parameter optimization corresponds to higher support for the “true” species tree. Indeed, inadequately assembled phylogenomic data might yield a strongly supported but incorrect tree (Lemmon et al. 2009), while lower support for the “true” topology might reflect a genuine biological pattern (e.g., incomplete lineage sorting or introgression, Degnan & Rosenberg 2009). Hence, optimizing assembly parameters to increase phylogenetic support might introduce biases toward certain topologies (Leaché et al. 2015b).

To further investigate these issues, we generated phylogenomic data sets of four European amphibian genera (*Discoglossus, Lissotriton, Rana, Triturus*), using double-digest RADseq (ddRADseq; Peterson et al. 2012) as well as Hybrid-Enrichment Capture (HE, based on the FrogCap probe set; Hutter et al. 2022). As the latter consists of genome-verified single-copy markers exempt of the biases of *de novo* clustering, we use it as a reference to evaluate the accuracy of inferences based on RADseq *de novo* assemblies produced using various CT values. Furthermore, the systematics and taxonomy of these genera are well known thanks to in-depth studies of their evolutionary histories and reproductive isolation, often based on contact zone analysis (e.g., Zieliński et al. 2013; Arntzen et al. 2014; Dufresnes et al. 2020a, 2020b). Therefore, it is possible to determine whether uncertainty in the results of species delimitation and phylogeny inferences from RADseq data could reflect genuine biological patterns or systematic biases introduced by inappropriate assembly parameters. Finally, the focal taxa differ drastically in genome size (c. 4-5 Gb in *Discoglossus* and *Rana*, c. 15-28 Gb in *Lissotriton* and *Triturus*; Gregory 2022), and the large genomes of the newts (*Lissotriton* and *Triturus*) can be expected to have an increased content of repetitive elements and longer introns (Smith et al. 2009). This could directly affect the efficiency of RADseq-based analyses by making ortholog clustering more challenging and decreasing the number of shared loci in the final assemblies (Andrews et al. 2016; Clugston et al. 2019).

Thus, the four focal amphibian genera of this study offer an ideal opportunity to investigate the impact of varying the CT on phylogenetic and MSC analyses, and to assess the extent to which assembly metrics accurately identify the optimal CT for a given RADseq data set in an empirical framework. To that end, we assemble the RADseq data sets using a wide range of CTs, and perform downstream concatenation-based phylogenetics, as well as MSC-based species-tree and species delimitation. In summary, the aims of the study are 1) to assess the impact of varying CT on downstream phylogenetic and MSC inferences by using HE data sets and previous contact zone analyses as references; 2) to identify assembly metrics that can be used to find the optimal CT for RADseq assembly; and 3) to integrate these metrics within a pipeline that can be readily used for future assemblies of RADseq data sets in the context of phylogenomic analyses.

## 2. Material and Methods

### 2.1 Sampling and RADseq laboratory work

The data used for this study was gathered in the context of a broader project focusing on several groups of European amphibians. We selected representatives from four genera (Fig. 1, Appendix 1): *Discoglossus* (N=26, all species included), *Lissotriton* (N=8, sampling focused on the *L. vulgaris*-*montandoni* complex), *Rana* (N=22, focused on the *R. temporaria* complex) and *Triturus* (N=18, focused on the Western European species).

**Figure 1.**
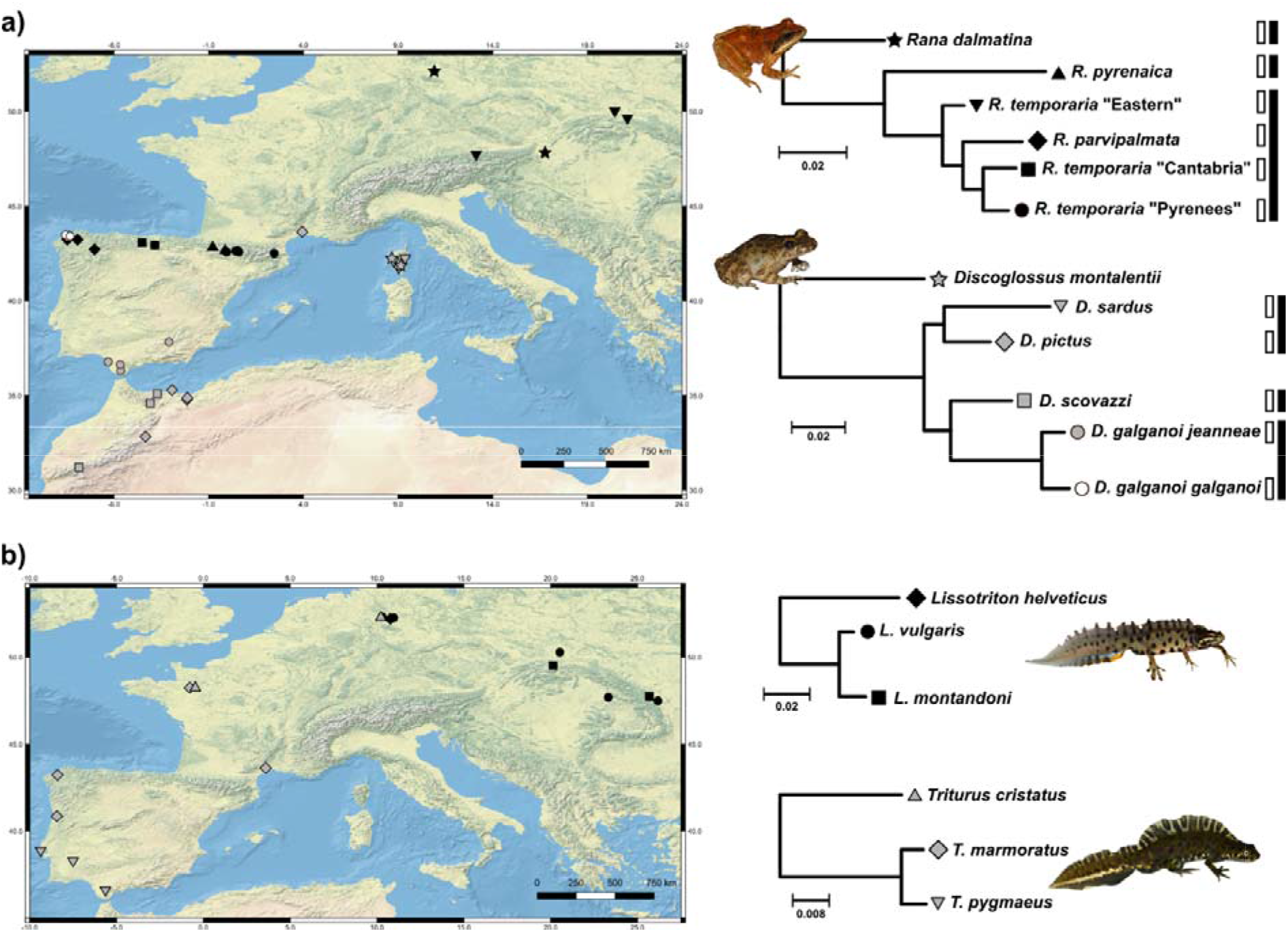
Simplified representations of the trees obtained from ML analyses of the concatenation of the HE loci in a) the anuran genera Rana and Discoglossus, and b) the urodelan genera Lissotriton and Triturus. All nodes received 100% aLRT support. Maps on the left show sampling sites. Complete trees are given in Figures S1LS4. Rectangles at the right of the Rana and Discoglossus trees show the partitions used to test the outcome of species delimitation analyses (open boxes = “split” scenarios, solid boxes = “lump” scenarios, cf. methods for more details).

Genomic DNA was extracted using the Macherey-Nagel NucleoSpin Tissue kit following the manufacturer’s instructions. We performed ddRADseq (Peterson et al. 2012) with library preparation as follows (per Recknagel et al., 2015 with modification of Illumina adapters; Rodríguez et al. 2017): 1 μg of DNA from each individual was double-digested using the PstI-HF and AclI restriction enzymes (NewEngland Biolabs); modified Illumina adapters with unique barcodes for each individual were ligated onto this fragmented DNA; samples were pooled at equimolarity; a PippinPrep was used to size select fragments around a tight range of 383 bp. Finally, enrichment PCR was performed to amplify the library using forward and reverse RAD primers. Sequencing was conducted on an Illumina Next-Seq at Glasgow Polyomics to generate 75 bp paired-end reads. After quality check using FastQC (www.bioinformatics.babraham.ac.uk/projects/fastqc/), raw reads were demultiplexed with stacks v2 (Rochette et al. 2019) and trimmed to 60 bp. Raw reads were submitted to the NCBI Short Read Archive under Bioproject <u>###### (to be added upon manuscript acceptance)</u>

### 2.2 Hybrid-Enrichment analyses

As a comparative basis for evaluating the quality of phylogenetic analyses performed with RADseq data, nuclear loci were also sampled through a Hybrid-Enrichment protocol, using a modified version of the FrogCap sequence capture probe set (Hutter et al. 2022). In brief, our goal was to create a set of universal amphibian markers that could be targeted using a single set of probes. We started by using the FrogCap markers (Hutter et al. 2022; https://github.com/chutter/FrogCap-Sequence-Capture) that were successfully captured broadly across Anura in the data from Hutter et al. (2022). Next, these markers were matched against the Salamander *Ambystoma mexicanum* genome (Keinath et al. 2018) and retained if they matched with ≥65% similarity. The total number of markers was 7,720 (Ultra Conserved Elements [UCEs]: 2,122; exons: 5,598). We further aimed to add additional universal markers that could potentially be compared and included in phylogenetic analyses with other groups of organisms. We selected the USCO (Universal Single Copy Orthologs) set of markers, which are orthologous genes found across deep evolutionary scales (i.e., Kingdom, Phylum, Class). To identify USCO markers present across Amphibia, we selected genomes of *Ambystoma mexicanum* and the frogs *Xenopus tropicalis*, *Rana catesbeiana*, *Nanorana parkeri*, *Rhinella marina*, and *Oophaga pumilio* (Sun et al. 2015; Hammond et al. 2017; Edwards et al. 2018; Keinath et al. 2018; Rogers et al. 2018). We used the program BUSCO v3.0.2 (Seppey et al. 2019) to locate and identify the Metazoa and Tetrapoda USCO markers in these genomes. We included all metazoan genes found in all the genomes, which totalled 677 out of 978 genes. We also added 29 genes from the Tetrapoda set to utilize remaining probes. Finally, to filter markers and create baits from the FrogCap markers, we followed Hutter et al. (2022; see Supplementary Material S1 of the present manuscript for more details). These steps resulted in a final set of 8,720 markers covering a total of 2,702,951 bp. The USCO markers totalled 14,153 baits (Metazoa: 13,196; Tetrapoda: 957) while the FrogCap markers totalled 25,884 baits (UCEs: 3,844; exons: 22,040).

Genomic DNA was extracted from the tissue samples using a PromegaTM Maxwell bead extraction robot and was quantified using a Promega QuantusTM fluorometer. Approximately 500 ng total DNA was acquired and set to a volume of 50 µl through dilution (with H20) or concentration (using a vacuum centrifuge) of the extraction when necessary. The genomic libraries for the samples were prepared by Arbor BioSciences library preparation service. Library preparation is detailed in the Supplementary Material S1.

Illumina sequence data were de-multiplexed using the Illumina software bcl2fastq. The bioinformatics pipeline for filtering adapter contamination, removing biological contamination, assembling contigs, and preparing sequence data for alignment is available at https://github.com/chutter/FrogCap-Sequence-Capture, and described in more detail in Supplementary Material S1. Next, the retained set of markers was aligned on a marker-by-marker basis using MAFFT local pair alignment (settings: max iterations=1000; ep=0.123; op=3; --adjust-direction; Katoh & Standley 2013). We screened each alignment for samples ≥40% divergent from consensus sequences, which were almost always incorrectly assigned to contigs. The obtained alignments were trimmed with trimAl (Capella-Gutiérrez et al. 2009), and subsequently realigned with MAFFT. Remaining poorly aligned regions were identified and trimmed using a custom R script. In brief, a consensus sequence was generated for each alignment, and the original sequences were divided in 80 bp long slices. The slices were then compared to the consensus sequence and removed when >40% of the positions differed. Resulting alignments that were <40 bp long or with less than 4 samples were filtered out. After trimming and filtering, the four data sets included 2,615 – 10,620 loci for a total of 921,033 – 6,751,735 bp (cf. Tab. S1 for more details).

The hybrid-enrichment data was used to generate reference phylogenetic trees in the four genera using both concatenated and species-tree analyses. First, the loci alignments were concatenated and used for maximum-likelihood (ML) phylogenetic inference in IQ-TREE v1.6.8 (Nguyen et al. 2015). The best-fitting substitution models and gene-partitions were selected with BIC in IQ-TREE’s implementation of ModelFinder (Chernomor et al. 2016; Kalyaanamoorthy et al. 2017). Branch support was assessed using the SH-like approximate likelihood ratio test (aLRT) with 1,000 pseudo-replicates. Second, single-gene trees were calculated in RAxML v8 (Stamatakis 2014) under a GTR+Γ substitution model and fed into ASTRAL II (Mirarab & Warnow 2015) for species-tree inference, an approach consistent with the MSC.

### 2.3 RADseq data assembly and parameter optimization

For each genus, RADseq *de novo* assemblies were generated from the demultiplexed reads using ipyrad v0.9.50 (Eaton & Overcast 2020), a software that processes RADseq data through seven sequential analytical steps (referred to later in this paragraph, see https://ipyrad.readthedocs.io/en/latest/7-outline.html for a detailed overview). In this pipeline, the CT is determined by a single parameter, the clustering threshold (#14 in the parameter file) which determines the percentage of similarity needed between two sequences to be clustered (i.e., considering sequences of 100 bp, a clustering threshold of 0.90 would cluster sequences with ≤10 substitutions). This parameter will be used at step 3 for intra-sample clustering of the reads (intra-sample Clustering Threshold, hereafter referred to as iCT) and at step 6 for between-samples clustering of the consensus sequences (between-samples Clustering Threshold, hereafter bCT). As this parameter only accepts a single value, ipyrad runs must be split into two parts to allow the iCT and bCT to differ. First, steps 1-5 of the analysis must be run with parameter #14 set to the iCT value. Secondly, the parameter file must be edited to change parameter #14 to the bCT value, and steps 6-7 must be run to complete the assembly (Paetzold et al. 2019; Karbstein et al. 2020; Hühn et al. 2022). We applied this strategy to find an “optimal” combination of iCT and bCT using the following approach: 1) the data sets were assembled with a range of iCT and a fixed bCT; 2) the resulting clusters and assemblies were examined to identify an “optimal” iCT (i.e., a value conservative enough to allow correct separation of orthologs, but not too high as to avoid under-merging of alleles at heterozygous loci, as detailed in the Results section) using both assembly metrics and downstream phylogenetic analyses; 3) the clusters generated with the “optimal” iCT were used as input to re-run steps 6 and 7 of ipyrad with a range of bCT values; 4) the resulting assemblies were investigated as in the second step to determine the “optimal” bCT. These steps are described in more detail in the following sections.

### 2.4 Intra-sample Clustering Threshold

For each of the four data sets, steps 2L5 of ipyrad were run with the following iCTs: 0.50, 0.60, 0.70 and 0.80-0.99 with an increment of 0.01. The *mindepth* parameter was set to 8 (based on Goudarzi et al. 2019; Rancilhac et al. 2019), and the remaining parameters were left as default. For each assembly, we used ipyrad s3 and s5 summary files to recover clusters-related metrics: total number of clusters, number of clusters passing the minimum depth filter (clusters_mindepth), depth of the clusters and number of clusters discarded due to high heterozygosity. Furthermore, to investigate the impact of the iCT on the loci assembled, steps 6 and 7 of ipyrad were run on the previously assembled clusters with a standard bCT of 0.90, which is ipyrad’s default and a commonly used value in phylogenetic studies (e.g., Eaton & Ree 2013; Paetzold et al. 2019). All the other parameters were left as default. The s7 summary files were used to collect loci-related metrics: number of loci in the final assembly, proportion of loci shared by ≥50% and ≥80% of the samples, total number of variable and parsimony-informative sites, and percentage of missing data in the concatenation and SNPs matrices. Custom scripts in R (R core team 2020) were used to visualize the impact of the iCT on these variables (all scripts are available at github.com/rancilhac/optiRADCT). Particularly, for each variable, a rate of change Δ was calculated as 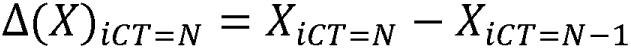.

Finally, the concatenated loci from each assembly were used to calculate ML phylogenetic trees using RAxML v8 under a GTR+Γ substitution model. The rapid bootstrapping algorithm was run with 100 replicates. The resulting trees were compared to the respective HE reference ML tree using two metrics: the Robinson-Foulds distance (RFD; Robinson & Foulds 1981) that measures topological difference only, and Kuhner & Felsenstein’s Branch Lengths Score (BLS; Kuhner & Felsenstein 1994), an extension of the previous that accounts for differences in both topology and branch lengths. It should be noted that four samples were missing from the HE datasets (three *Discoglossus* and 1 *Triturus*), and one from the RADseq datasets (Appendix 1). These samples were removed to compare the trees. Distances were computed using custom R scripts based on ape v5.6-2 (Paradis et al. 2004), which was also used to assess changes in phylogenetic uncertainty (i.e., bootstrap support of the branches [BS]) and branch lengths, depending on the iCT. To summarize branch lengths into a single variable, we calculated changes of the tree diameter as defined by Mai & Mirarab (2018), which is the longest pairwise patristic distance between tips of the tree. Pairwise distances were calculated using the package adephylo v1.1-11 (Jombart & Dray 2010).

### 2.5 Between-sample Clustering Threshold

The results of the previous analyses were used to identify a threshold above which under-merging of heterozygous loci was pervasive, resulting in a sharp increase of the number of clusters and missing data in the final matrices, and a strong decrease of the depth of the clusters, number of loci and informativeness of the assembled matrices (the underlying rationale is detailed in the Results and Discussion sections). The “optimal” iCT was selected as the highest value under this threshold, to ensure correct splitting of paralogs while avoiding under-merging of alleles from orthologous loci. The clusters assembled with the “optimal” iCT value were used to run steps 6L7 of ipyrad with the following bCTs: 0.50, 0.60, 0.70 and 0.80L0.99 with an increment of 0.01. Other parameters were left as default. Variation in the assembly metrics depending on the bCT was investigated using the same criteria as for the iCT.

### 2.6 Impact of the bCT on phylogenetic and Multi-Species Coalescent analyses

For the 92 assemblies, the concatenated loci were used to calculate ML phylogenetic trees with RAxML, using the same settings as before. The quartet-based method tetrad (https://github.com/ eaton-lab/tetrad) was also used to infer species trees from the unlinked SNP matrices outputted by ipyrad (i.e., the best-covered SNP is selected from each locus or taken as random in case of equality). The impact of the bCT on the topologies, resolution and branch lengths of the inferred trees was investigated with the same approach as the iCT. As tetrad does not compute branch lengths, only topologies and supports were compared in this case.

For the *Discoglossus* and *Rana* data sets, we conducted further MSC-based inference of species tree and species limits using the SNAPP/BFD* framework (Bryant et al. 2012; Leaché et al. 2014) implemented in BEAST 2 (Bouckaert et al. 2019). Because this approach is very sensitive to missing data (Schmidt-Lebuhn et al. 2017), unlinked SNP matrices were filtered to keep only sites covering all samples. Prior to that, individuals with missing data at >85% of all SNPs were removed to mitigate the data loss. In the *Discoglossus* data set, this resulted in the exclusion of *D. montalentii* as an effect of loci dropout but including it would have drastically reduced the SNP matrices, making it difficult to determine whether uncertainty was due to an improper bCT or the low amount of data. First, SNAPP was run seven times for each genus, using SNP matrices assembled with the “optimal” bCT as identified in the previous steps (0.91 in both cases, cf. Results), and six alternative bCTs representing various levels of over-merging (0.80, 0.85 and 0.88) and under-merging (0.93, 0.95 and 0.99). In total, 14 SNP sets were analysed (Tab. S2). The priors were set following Leaché & Bouckaert (2018), and the analyses were run for a maximum of 15,000,000 steps sampling every 10,000. After the first 5,000,000 steps, convergence was checked with Tracer v1.7 (Rambaut et al. 2018). Analyses were stopped if they showed signs of adequate convergence (i.e., Effective Sampling Size of the parameters >200 and no clear trend in the posterior trace) or left to run until convergence was reached. For each SNAPP run, the posterior sampling of trees was visualized in Densitree v2.2.7 (Bouckaert 2010). Furthermore, variation in the heights and posterior probabilities of the nodes, and in the population size estimates (θ) in terminal branches depending on the bCT was also assessed. Second, the BFD* method was applied to compare three alternative species delimitation scenarios in each genus (Fig. 1a): the current species-level taxonomy, a “split” scenario (some conspecific populations are split into separate species) and a “lump” scenario (some distinct species are considered conspecific). The marginal-likelihoods (Pr) of SNAPP runs were calculated using the Path-sampling algorithm implemented in BEAST 2, which was run with 48 iterations of 400,000 steps. For each SNP matrix, the alternative scenarios were compared to the reference taxonomy using a Bayes Factor (BF) score which was calculated as follow: 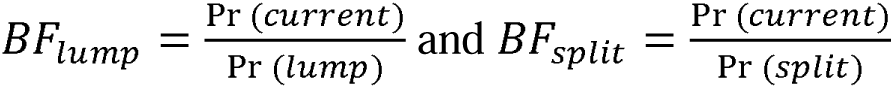.

## 3. Results

### 3.1 Hybrid-Enrichment-based phylogenetics

In all four genera, concatenation and species-tree approaches applied to the HE data yielded concordant and fully resolved phylogenetic trees (Fig. 1, Figs. S1L4; ASTRAL species trees are not illustrated as their topologies were identical to concatenation trees), that are mostly concordant with published phylogenetic studies of these taxa. Although systematics is not the scope of the present study, it is worthwhile to note that *Rana temporaria* is divided into three deep lineages that are paraphyletic respective to *R. parvipalmata* (Fig. 1a, Fig. S3). This pattern will not be further analyzed here but points to the need of future studies to clarify the species limits and phylogeographic history of this genetically diverse group.

### 3.2 RADseq data: effect of the intra-sample Clustering Threshold

For this first step, 23 assemblies were generated for each data set. Loci sequences and SNP matrices outputted by ipyrad, as well as spreadsheets providing detailed metrics for each assembly, have been uploaded to https://zenodo.org/record/7829243 (DOI: 10.5281/zenodo.7829243). In all cases, the number of constructed clusters increased, and their depth decreased, with increasing iCT (Fig. S5). Both variables showed a break at iCT=0.95 in all four data sets, above which the variation generated by each increment of the iCT was substantially higher. Finally, the number of clusters rejected by the heterozygosity filter followed a plateau for iCTs between 0.80 and 0.91 before decreasing drastically and stabilizing near zero for iCT > 0.95 (Fig. S5).

Subsequently assembling these clusters with a standard bCT of 0.90 revealed substantial variation in the resulting data sets. The total number of loci recovered regularly increased until reaching a maximum around iCT=0.90, after which it decreased steeply. This general trend is also found when focusing on the proportion of loci shared by ≥50% and ≥80% of the samples (Fig. 2a). Particularly, while the proportion of the “shared loci” relative to the total number of loci is rather stable for iCTs ranging between 0.50 and 0.90, it starts decreasing between 0.90 and 0.95, and drops steeply past the latter value. The same pattern was also observed for the number of variable and parsimony-informative sites (Fig. 2b). Finally, the proportion of missing data in the SNP (31.6% – 66.1%) and concatenation (34.5% – 71.8%) matrices was relatively stable for iCTs<0.90, above which it increased drastically (Fig. 2). All data sets were largely concordant in supporting two distinct breaks in the rate of change of these variables, for values of iCT=0.90 and 0.95, although the trend was somewhat weaker for *Triturus* (Fig. 2). The first of these breaks was not very strong, but iCT=0.95 marked a threshold above which each increment of the iCT generated a strong drop in the number of shared loci and variable and parsimony informative sites, as well as a strong increase in the proportion of missing data.

**Figure 2.**
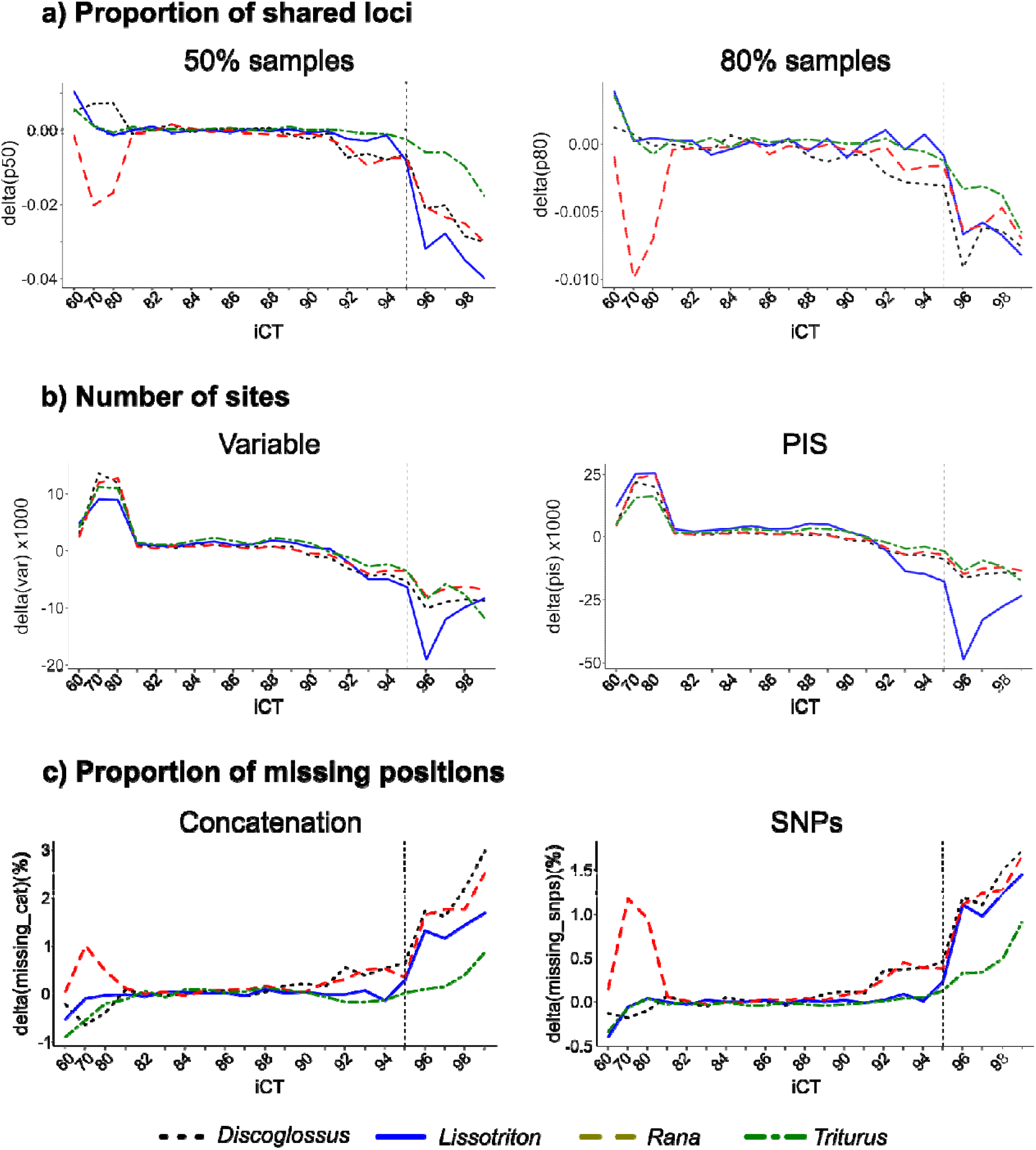
Impact of the intra-sample Clustering Threshold (iCT) on RADseq assemblies with a standard between-samples Clustering Threshold (bCT)=0.90. a) variation in the proportion of assembled loci shared by 50% and 80% of the samples. b) variation in the number of variable and parsimony informative sites (PIS) in the final matrices. c) variation in the percentage of missing data in the concatenation and SNP matrices. The graphs show the rate of change Δ of the variables when increasing the iCT. Vertical dashed lines show iCT=0.95, which marks a break in the variation of Δ in most cases (but see Discussion). Note: the scale of the y-axis is not proportional to improve the visualisation of patterns at high iCTs.

Despite some dissimilarities, the concatenation-based phylogenetic analyses yielded overall consistent topologies across iCT values. All inferred trees in Newick parenthetical format are available at https://zenodo.org/record/7829243. In all genera, inter-specific relationships were stable and fully supported (i.e., BS=100) for all iCT values, even unrealistically low (e.g., 0.50, 0.60) or high (e.g., 0.99) ones. These relationships were identical to those of the HE reference trees, except for *Rana* in which the position of *R. parvipalmata* differed (sister to all Spanish *R. temporaria* in the HE trees vs. sister to the Cantabrian populations only in the RADseq trees). HE and RADseq trees differed slightly in intra-specific relationships in all genera except *Lissotriton*, resulting in Robinson-Foulds Distances (RFD) >0 (Fig. 3a). Although somewhat lower bootstrap values were observed mostly at higher iCT values >90 (Fig. 3b), the topological variation was apparently not driven by the iCT (Fig. 3a), and likely reflects genuine phylogenetic uncertainty at the intra-specific level, where strictly bifurcating trees might not accurately represent the relationships between populations. Accordingly, mean BS values were >90 in all trees (Fig. 3b) and their variation did not appear to be matched to changes in the iCT. Finally, increasing the iCT resulted in a slight but steady increase of the tree diameter (Fig. 3c), which translated into a decrease of the Branch Length Scores (not illustrated).

**Figure 3.**
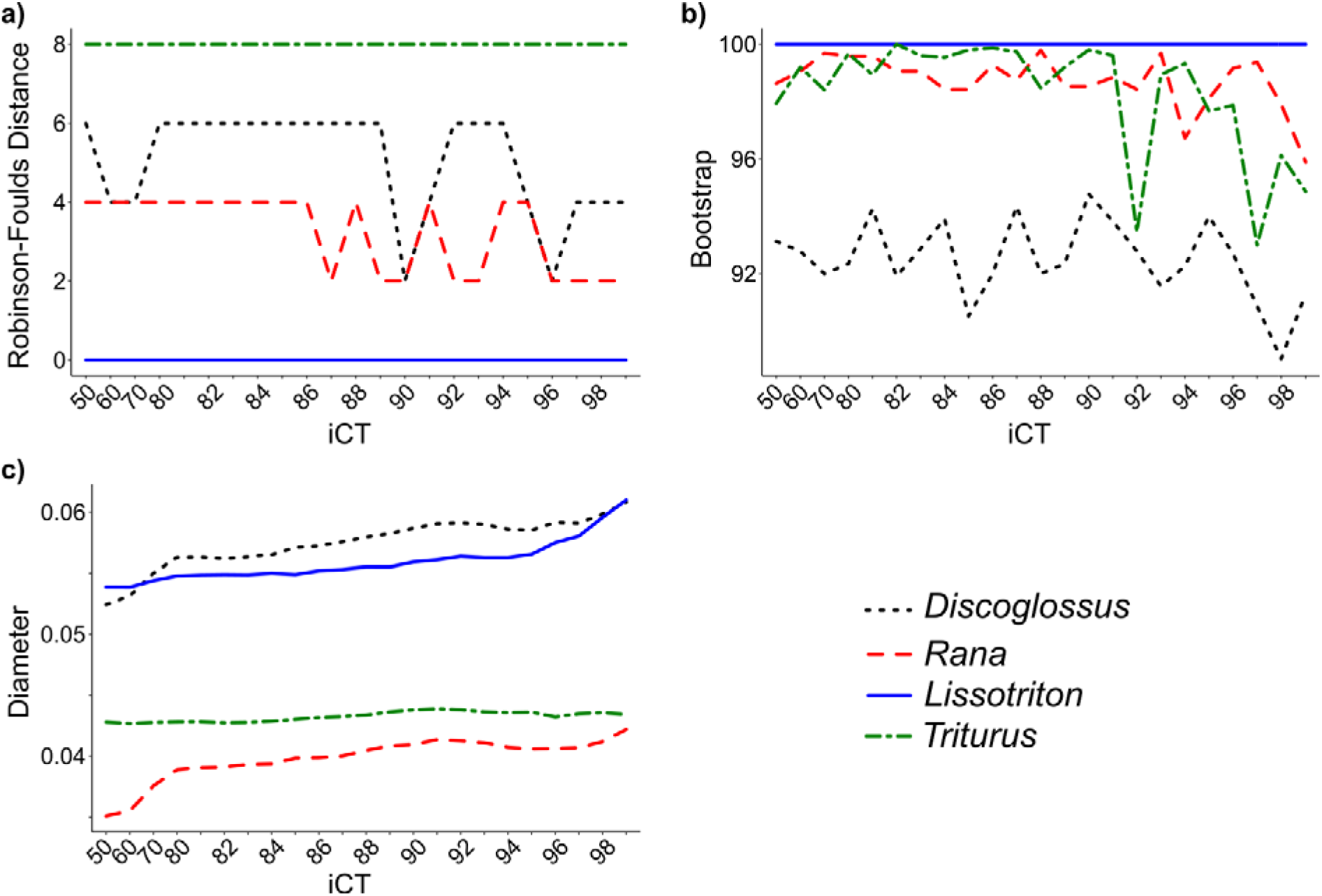
Impact of the intra-sample Clustering Threshold (iCT) on concatenation-based phylogenetic reconstructions, represented as the change of three metrics as a function of the iCT: a) Robinson-Foulds Distance to HE reference trees; b) mean bootstrap support of the RADseq trees; c) diameter of the RADseq trees. Note: the scale of the y-axis is non-proportional for improved visualisation of patterns at high iCTs.

### 3.3 RADseq data: impact of the between-sample Clustering Threshold

In the four genera, clusters- and loci-related assembly metrics were relatively stable for 0.80<iCT< 0.90 (Fig. 2, Fig. S5). Especially, the change of three clusters-related metrics – the number of clusters, depth of the clusters and number of clusters of high heterozygosity – showed a break at iCT=0.95 in all four genera. Loci-related metrics, on the other hand, were less straightforward to interpret, particularly because *Triturus* showed less variation than the three other taxa. While some signs of under-merging were noticeable from iCT=0.90 (e.g., increase of missing data in *Discoglossus* and *Rana*), we find that the most striking decrease of shared loci and variable sites, and increase of missing data, occur for iCT>0.95 in all four taxa. We interpret this pattern as indicating that iCT=0.95 represent a threshold above which under-merging of heterozygous loci becomes pervasive (cf. Discussion for more details). Hence, we considered this value as the “optimal” iCT for the next analytical steps. The clusters constructed with iCT=0.95 were assembled into loci with 23 bCTs ranging from 0.50 to 0.99.

The effect of the bCT on the informativeness and completeness of the assemblies was overall similar to that of the iCT (Fig. 4; assemblies and metrics are available at https://zenodo.org/record/7829243). Trends in loci-related metrics identified two bCT values inducing noticeable breaks. In the one hand, the proportion of loci shared by ≥50% of the samples and proportions of missing data in concatenation and SNP matrices remained relatively constant for 0.80<bCT<0.91, above which they decreased consistently in all four genera. On the other hand, the number of variable sites and PIS decreased clearly only for bCT >0.95. The pattern of change in the variation of the proportion of loci shared by ≥80% of the samples was not consistent across genera, as this metric decreased only for bCT >0.95 in *Triturus*, vs. bCT >0.91 in the three other genera, although with different levels of amplitude.

**Figure 4.**
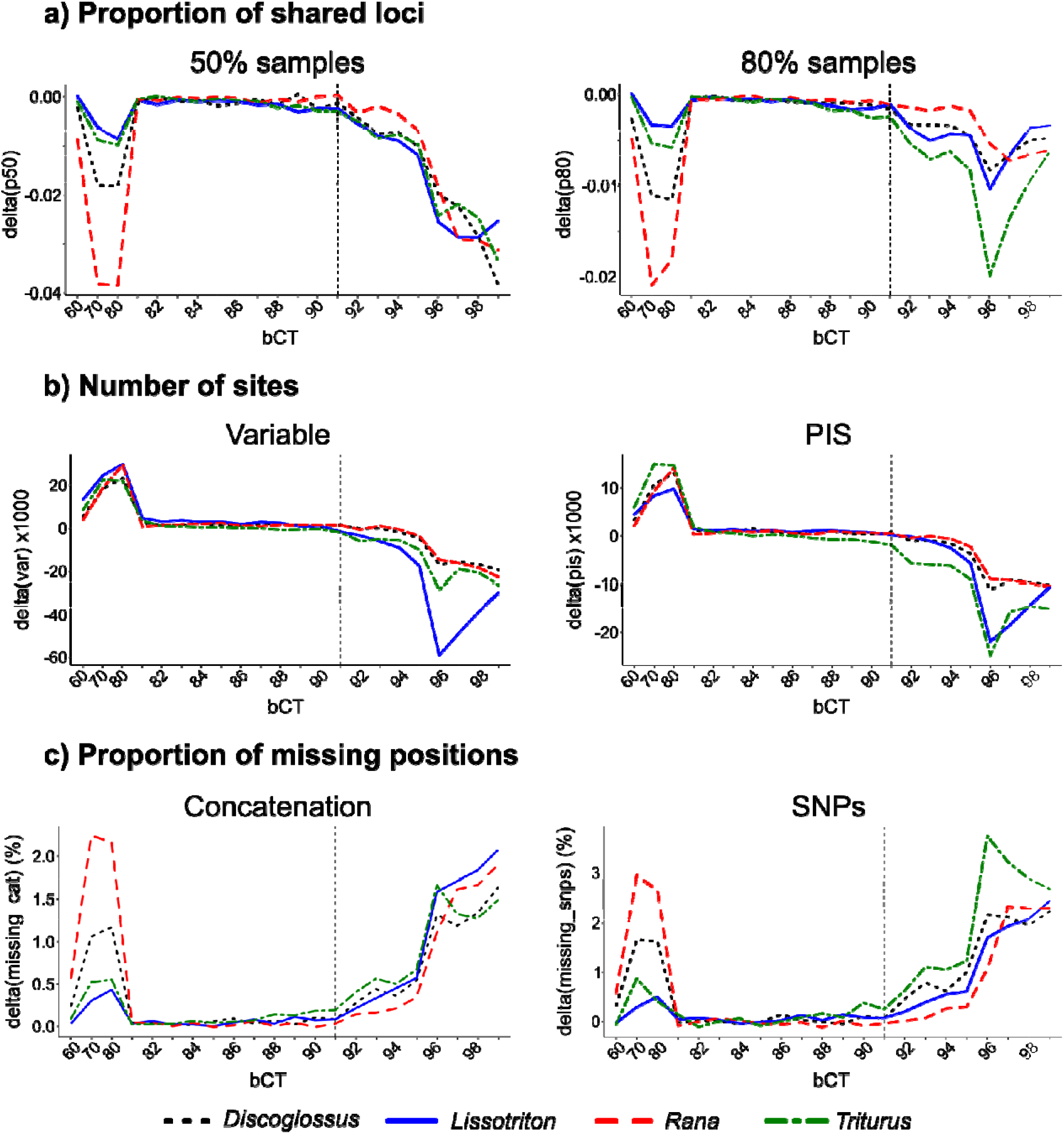
Impact of the between-sample Clustering Threshold (bCT) on RADseq assemblies with an iCT=0.95. a) variation of the proportion of assembled loci shared by 50% and 80% of the samples. b) Number of variable and parsimony informative sites in the final matrices. c) Proportion of missing data in the concatenation and SNP matrices. The graphs show the rate of change Δ of the variables when increasing the bCT. Vertical dashed lines show bCT=0.91, which marks a break in the variation of Δ. Note: the scale of the y-axis is non-proportional for improved visualisation of patterns at high bCTs.

The extent of the variation induced by changes of the bCT was greater than for the iCT. For example, increasing the bCT from 0.91 to 0.99 resulted in an average 4.70% decrease of the proportion of loci shared by 80% of the samples, and an average 11.6% increase of missing data in the SNP matrices. Comparatively, the same change of iCT induced a decrease of 2.9% and an increase of 6.8% in these two metrics, respectively.

As for the iCT, concatenation-based phylogenetic analyses were resilient to changes in the bCT, and all assemblies yielded the same topologies as at the previous step with high support for inter-specific relationships (all inferred trees are available in Newick parenthetical format at https://zenodo.org/record/7829243). In *Discoglossus*, the BS of two inter-specific branches decreased to minima of 76 and 87, respectively, for bCT≥0.95, whereas bCTs≤0.60 failed to support the monophyly of *D. g. galganoi* (BS≤59). In *Rana*, the BS of the branch separating *R. parvipalmata* and its closest relative decreased to a minimum of 87 for bCTs≥0.98. Otherwise, all bCT values recovered species monophyly as well as inter-specific relationships with maximum support in all four genera. Consequently, RF distances to the reference HE trees did not change as a function of the bCT (Fig. 5a). Similarly, variation in BS values was not associated with the bCT in a clear way (Fig. 5c). In accordance with the observed topological stability, the BS was overall high, with few outliers highlighting phylogenetic uncertainty at the intra-specific level, regardless of the bCT. However, in *Discoglossus* and *Triturus*, bCTs ≤80 and ≥95 seemingly yielded overall lower BS.

**Figure 5.**
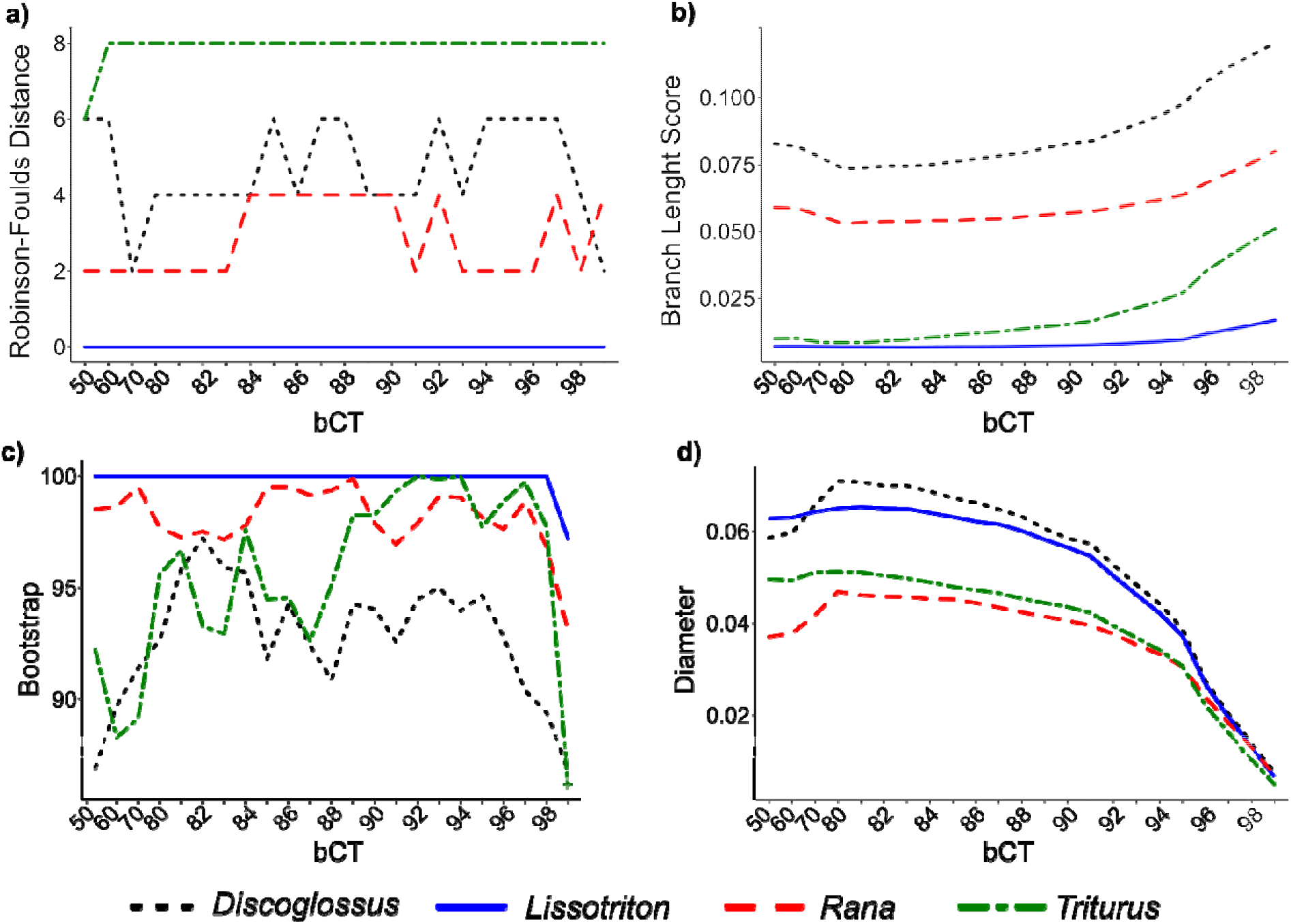
Impact of the between-sample Clustering Threshold (bCT) on concatenation-based phylogenetic reconstructions, represented as the change of four metrics as a function of the bCT: a) Robinson-Foulds Distance to HE reference trees; b) Branch Length Score to HE reference trees; c) mean bootstrap support of the RADseq trees; d) diameter of the RADseq trees. Note: the scale of the y-axis is non-proportional for improved visualisation of patterns at high bCTs.

The most striking effect of the bCT was on the branch lengths (Fig. 5d). For bCTs ≥0.80, each increment resulted in a contraction of the branches, which was particularly important for bCT >91. Increasing the bCT from 0.91 to 0.99 resulted in a dramatic contraction of the tree diameters by an average factor of 7.36 (5.57 – 8.42). This pattern was reflected by the BLS between RADseq and HE trees (Fig. 5b), which increased steadily with increasing bCTs, particularly for bCT >0.91. To further investigate the behaviour of branch lengths in concatenation trees when increasing the bCT, we plotted the trees’ diameters depending on the number of PIS (Fig. 6a) and the proportion of missing data (Fig. 6b). In both cases, we found that the association between both variables changed across bCTs. Indeed, for high bCTs, the diameter seems to increase linearly according to the number of PIS and decrease linearly depending on the proportion of missing data. However, for bCTs between 0.80 and c. 0.90, the diameter’s increase continues although the number of PIS decreases, and the proportion of missing data remains stable. The bCT value marking the transition between these two trends is variable depending on the taxon considered, ranging from 0.90 to 0.95. In all four cases, the data sets assembled with bCTs of 0.50, 0.60 and 0.70 were outliers. Finally, to determine whether the branch length contraction was proportional across branches within a single tree, we normalized the lengths of internal branches (which were shared across all trees) by dividing them with the respective tree’s diameter. In case the decrease in branch length affects all branches proportionally, we expect that normalized branch lengths would remain constant. However, we observe that normalized branch lengths vary, following trends that are branch-dependent (Fig. 6c). The longest branches show the strongest proportional contraction, while shorter branches become proportionally longer, or remain stable.

**Figure 6.**
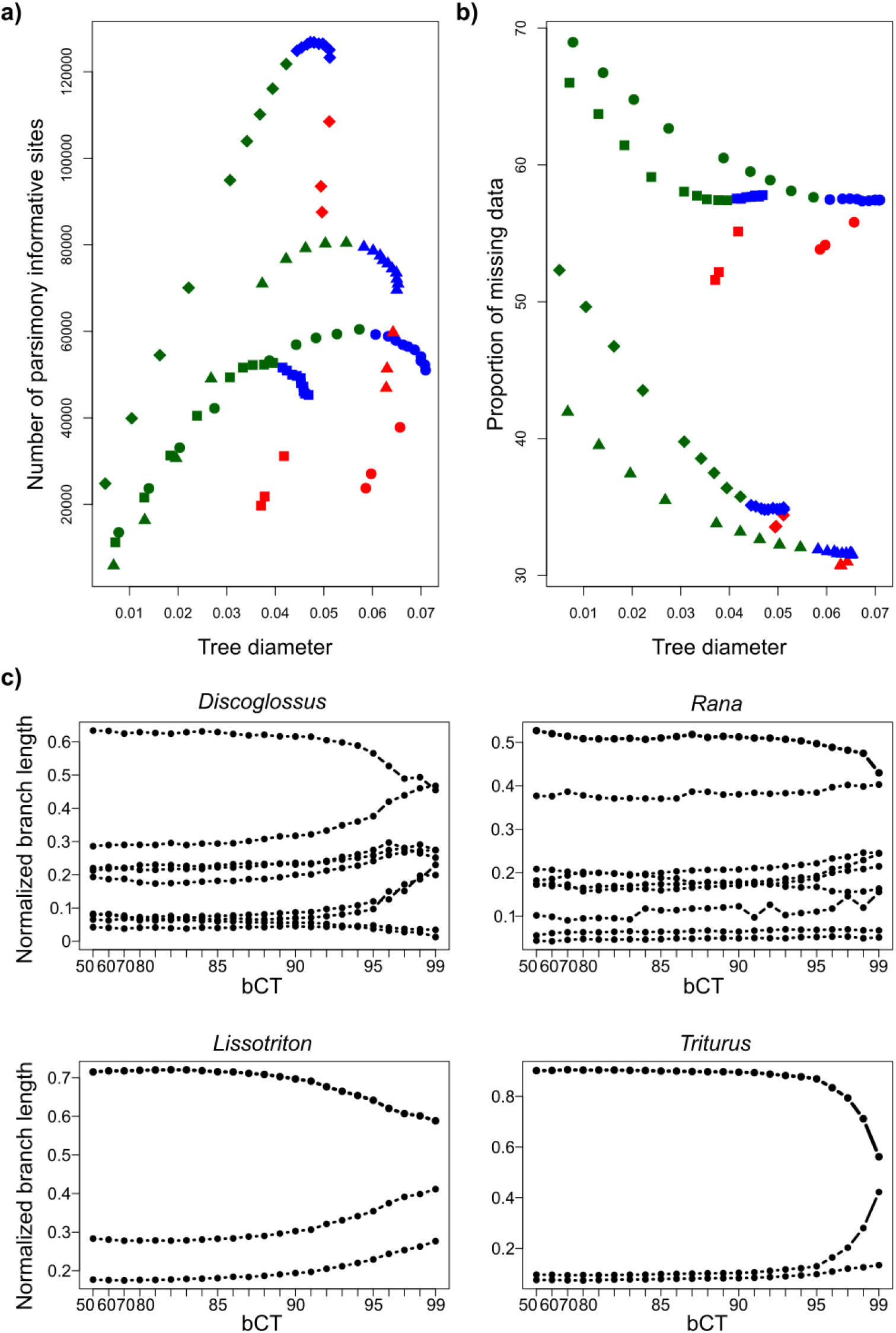
Details of the impact of the between-sample Clustering Threshold (bCT) on branch lengths of concatenation trees. a) Relationship between the trees’ diameters and the number of parsimony informative sites. Dots = Discoglossus; squares = Rana; triangles = Lissotriton; diamonds = Triturus. b) Relationship between the trees’ diameters and the proportion of missing data in the assemblies. In these two graphs, the colors of the points denote the bCT used: red = bCT ≤ 0.70; blue = 0.90 ≥ bCT ≥ 0.80; green = bCT ≥ 0.90. c) Change of normalized length (i.e., divided by the tree diameter) of internal branches when increasing the bCT. Each line represents a given branch, and each dot the length of this branch for a given bCT. Note: the scale of the y-axis is non-proportional for improved visualisation of patterns at high bCTs.

Finally, quartet-based species trees were reconstructed using Tetrad. As for the concatenation analyses, both the topologies and the branch supports were remarkably resilient to changes in the bCT (Fig. S6), although bootstrap supports were overall lower. The inferred species-level relationships were identical to those of concatenation trees inferred from both HE and RADseq data. In the case of *Rana*, the position of *R. parvipalmata* was identical to the HE reference topology (i.e. sister to all Spanish *R. temporaria*), although with low support.

### 3.4 RADseq data: impact of the bCT on Multi-Species Coalescent analyses

Species trees inferred with SNAPP were generally resilient to changes in the bCT (Fig. 7). In the case of *Rana* (Fig. 7a), the “optimal” bCT (0.91) and two “sub-optimal” values (0.88 and 0.93, respectively) recovered a topology identical to that of the HE reference tree, with most nodes fully supported (i.e., Posterior Probability [PP]=1.00). Three alternative topologies were recovered with other bCT values, that differed in the relationships among *R. parvipalmata* and *R. temporaria* lineages. Increasing the bCT did not induce a clear change in the PP of the nodes. In *Discoglossus*, all analyses converged to a single topology (Fig. 7b), that differs from those previously obtained by placing *D. scovazzi* as sister to the *D. pictus* / *D. sardus* clade rather than to *D. galganoi*. This topology was well supported, except for bCT=0.99, where all nodes had a posterior probability <0.65. One clear effect of increasing the bCT on SNAPP species trees was a decrease of node heights, especially the roots, as exemplified by analyses for bCT=0.99 (Fig. 7a-b, Fig. S7). On the contrary, θ estimates were negligibly affected by changes to bCT (Fig. S8). In some lineages, higher bCT values yielded either higher or lower estimates, but no clear trend could be observed, and large confidence intervals made interpretation of the results inconclusive. Finally, the results of species-delimitation analyses remained constant across bCT values (Fig. 7c): in all cases, models over-estimating the species richness were preferred to both the accepted taxonomy and models under-estimating the species richness.

**Figure 7.**
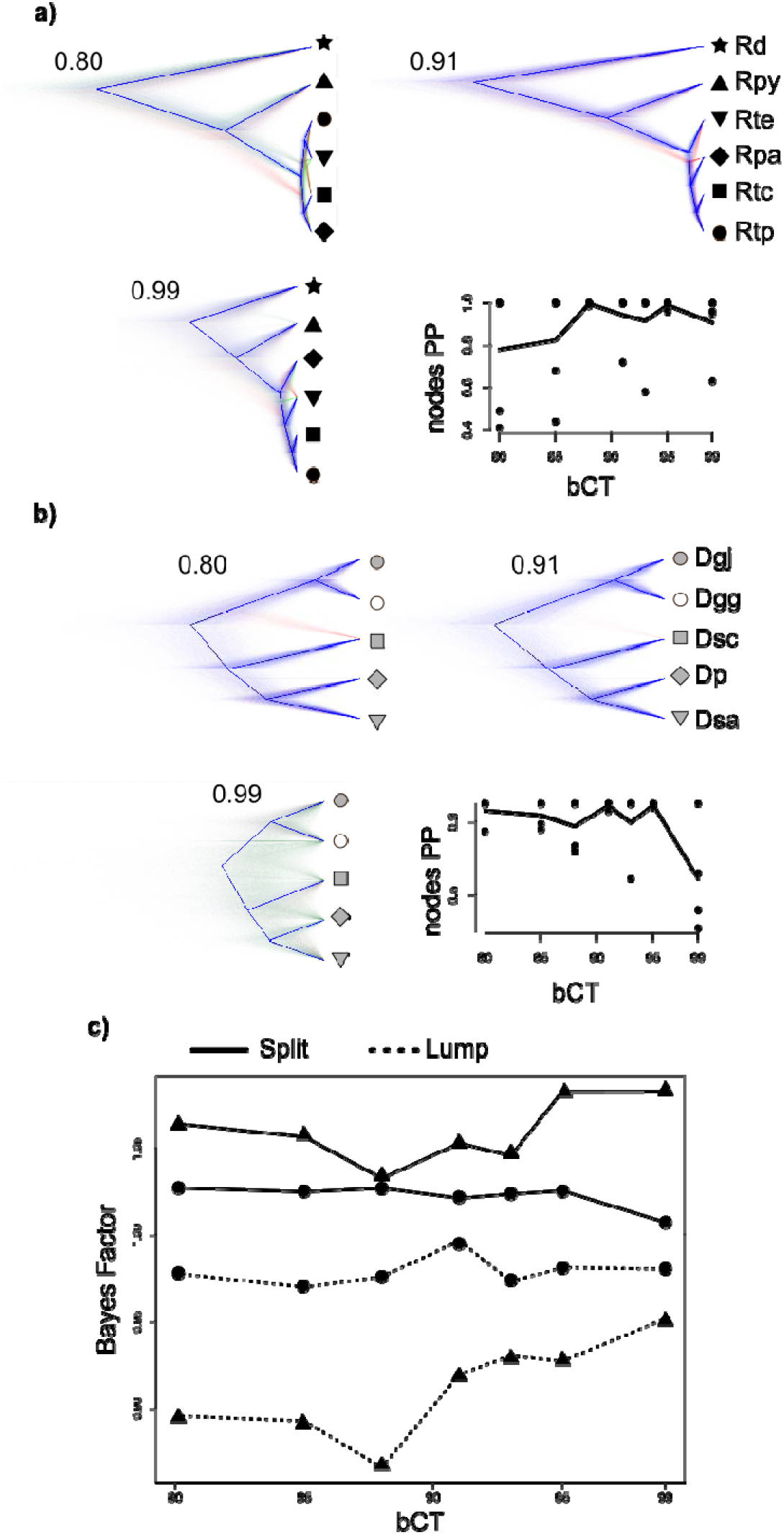
Impact of the between-samples Clustering Threshold (bCT) on Multi-Species Coalescent analyses. Cloudograms of the posterior distribution of SNAPP species trees in a) Rana (Rd=R. dalmatina, Rpy=R. pyrenaica, Rte=R. temporaria “Eastern”, Rpa=R. parvipalmata, Rtc=R. temporaria “Cantabria”, Rtp=R. temporaria “Pyrenees”) and b) Discoglossus (Dgj=D. galganoi jeanneae, Dgg=D. g. galganoi, Dsc=D. scovazzi, Dp=D. pictus, Dsa=D. sardus) for bCTs of 0.80, 0.91 and 0.99, with different colors highlighting competing topologies, and thick blue lines showing the consensus topology. The graphs show node support (posterior probability [PP]) as a function of the bCT (solid line = mean PP). The symbols at the tips refer to those used in Figure 1. c) Species delimitation analyses in BFD* in both Rana (dots) and Discoglossus (triangles). In both cases, the accepted taxonomy is compared to “split” (more species) and “lump” (less species) scenarios. The species hypothesis corresponding to each scenario is given in Figure 1a.

## 4. Discussion

In this study, we explore the effects of varying intra-and between-samples clustering thresholds for *de novo* assembly of multispecies RADseq data sets on downstream analyses widely used in systematics studies. Although our results are applicable to studies using RADseq markers at any scale, they perhaps bear relevance for studies with limitations on number of individuals sequenced per species, such as is typical for phylogenomic, species delimitation, systematic and taxonomic endeavours.

### 4.1 Assembly-metrics based identification of the optimal Clustering Thresholds in RADseq phylogenomic studies

It is common practice in RADseq-based phylogenetic studies to perform both intra- and between-samples clustering steps using the same Clustering Threshold (but see Hühn et al. 2022; Karbstein et al. 2020; Paetzold et al. 2019 for recent examples of differential optimisation). As the divergence between alleles within a single individual is expected to be substantially lower than between distinct species (with the exception of introgressed alleles), we here explored the effects of both intra-sample (iCT) and between-samples (bCT) Clustering Thresholds independently. When varying the former, we found that the properties of both clusters and loci were affected. Indeed, increasing the iCT yielded 1) a higher number of clusters, but of lower depth and heterozygosity, and 2) a decrease in shared loci across samples and overall loci variability, and an increase in the proportion of missing data. These changes reflect the “under-merging phenomenon” (McCartney-Melstad et al. 2019; Hühn et al. 2022), where an increase of the iCT will ultimately lead to splitting alleles from heterozygous loci into independent clusters of lesser depth, while lower thresholds can lead to the clustering of non-orthologous sequences, creating fewer clusters of higher depth (Paris et al. 2017). This will in turn affect the properties of the assembled loci because a higher iCT will increase the number of clusters discarded due to low read coverage, and artificially increase the number of invariant loci. In this context, proper optimization of the iCT appears crucial for *de novo* assembly of multispecies RADseq data sets. Hence, it is of interest to note that a noticeable break in the rate of change of most clusters- and loci-related metrics could be identified for iCT>0.95, although trajectories varied depending on the metrics and taxa considered. For example, the number of clusters above minimum depth did not show a clear break when increasing the iCT, likely because the number of clusters being lost to the depth filter for high iCTs was compensated by the overall increase of the number of clusters. Regarding the loci-related metrics, the increase of missing data in *Discoglossus* and *Rana* at lower iCTs compared to the two newt genera could be due to their deeper crown age and the higher number of samples in these data sets, which could accentuate the effect of allelic dropout. The attenuated variation in most loci-related metrics in the *Triturus* data set remains an unexplained idiosyncrasy; given that no such pattern was observed in the second newt genus in our analysis (*Lissotriton*), the unusually large genome size of these urodeles cannot be invoked as an explanation for this pattern.

Because of the variation in trajectories of these metrics, it seems unrealistic to define a simple and accurate heuristic solution to the clustering threshold optimization problem. However, we argue that the breaks identified at iCT=0.95 are informative in this context. Indeed, iCT values >0.95 provided the most noticeable and consistent signs of under-merging, namely a sharp increase in the number of clusters, a sharp decrease of their depth and heterozygosity, a decrease in the proportion of shared loci and variable sites, and an increase in the proportion of missing data. Hence, iCT=0.95 can be used as an “under-merging threshold” above which the partitioning of heterozygous loci becomes highly likely (similar to the “transition area” defined by Hühn et al. 2022). On the other hand, it is difficult to determine exactly how low the iCT can be set without the clustering of paralogs becoming a problem. An approach that is often employed is to maximize the number of PIS in the final assembly, corresponding to iCTs of 0.89 – 0.91 in our data sets. As this level of divergence is closer to what would be expected among species rather than at the intra-specific level, we advocate that this high number of PIS likely reflects the clustering of paralogs rather than a correct reconstruction of genetic variation within the data sets. Hence, we decided to select an iCT as conservative as possible while remaining below the “under-merging threshold” (i.e., 0.95), as a trade-off between avoiding paralog over-merging and ensuring that the level of under-merging remains low.

Next, we used the clusters yielded by the “optimal” iCT to produce loci assemblies under a range of bCTs. As expected, we found that increasing the bCT resulted in a lower proportion of shared loci, a lower number of variable sites, and an increase in the proportion of missing data, concordantly in the four studied taxa. This similarity suggests that, when increasing both iCT and bCT, under-merging of the loci across samples affects the assemblies in a similar way as under-merging of alleles within the samples. However, contrary to the pattern observed with the iCT, changes in these three metrics did not indicate a single “under-merging threshold”. Indeed, while some metrics (proportions of loci by ≥80% of the samples, number of variable sites and PIS) showed a threshold at bCT=0.95, the other metrics show sharp changes for bCT>0.91, while being stable for 0.80≤bCT≤0.91. Loci under-merging is also a likely explanation for this pattern, as it would result in assembling more loci with lower individual coverage. Thus, it seems that under-merging affects the assemblies even for bCT <0.95, although not all loci-related metrics supported it.

Interestingly, bCT=0.91 also produced the most informative assemblies in all taxa but *Triturus*, maximizing the number of variable sites and PIS. However, as discussed in more detail below, the effect of the iCT and bCT on phylogenetic inference was low, and thus properties of inferred trees could not be used as decisive optimization criteria. The only exception were branch lengths in concatenation trees, which consistently decreased with increasing bCTs. Determining which bCT value produces the most accurate estimation of branch lengths appears difficult, as the true values are not known, and branch lengths in HE reference trees are expected to differ due to different mutation rates (especially in coding regions). Nevertheless, we observe that branch contraction was much accentuated for bCT>0.91, concordantly with several loci-related metrics, as stated above. Branch contraction is expected to result from reduced informativeness of the assemblies and should thus be an indicator of under-merging of loci. Consequently, following the same reasoning as for the iCT, we consider bCT=0.91 as an “under-merging threshold”, above which heterozygous loci are likely to be split, and select it as the “optimal” value for our data sets.

In summary, we have shown that the trajectories of clusters- and loci-related metrics were quite variable, and somewhat taxa-dependent, when varying the iCT and bCT, making the optimization of these parameters a non-trivial problem. However, we could identify thresholds for both parameters above which it is safe to assume that clusters and loci under-merging becomes pervasive. In the absence of clear evidence for “optimal” CT values, we conservatively propose to use the highest CT below the “under-merging threshold”. Doing so, we could identify iCT and bCT values that yielded good quality data sets (i.e., supporting well resolved trees concordant with reference topologies, in both concatenation and species tree analyses, and mitigating branch contraction), following a two-step approach. We thus propose the following decision framework to help optimize these parameters in future RADseq-based studies that include samples of different species-level lineages (summarized in Tab. 1):

**Table 1.** Optimization framework for the intra-sample (iCT) and between-sample (bCT) Clustering Thresholds. Depending on whether the Clustering Threshold is set too low (over-merging) or too high (under-merging), the effects on clusters, loci and downstream phylogenetic analyses are described (Diagnosis column) and guidelines are proposed for optimization (Optimization solutions column). In the Diagnosis column, arrows describe metrics trajectory. One arrow means moderate variation, two arrows strong variation. The Observed values column gives the Clustering Threshold values corresponding to each case in our four datasets.

| Parameter | Clustering scenario | Observed values | Diagnosis |  |  | Optimization solutions |
| --- | --- | --- | --- | --- | --- | --- |
|  |  |  | Clusters | Loci | Phylogenetic trees |  |
| iCT | Over-merging | 0.50 – 0.94 | ↓ N clusters<br>↑ depth<br>↑ clusters rejected by heterozygosity filter | Loci metrics remain stable | No clear effect | Use the highest iCT below the under-merging threshold (strong break in the change of clusters and loci metrics) |
|  | Under-merging | 0.96 – 0.99 | ↑↑ N clusters<br>↓ depth<br>↓↓ clusters rejected by heterozygosity filter | ↓↓ proportion of shared loci and variable sites<br>↑↑ proportion of missing data | No clear effect |  |
| bCT | Over-merging | 0.50 – 0.90 | NA | Loci metrics remain stable | ↓ bootstrap support for very low bCTs (low effect) | Use bCT that maximizes informativeness while below the under-merging threshold (strong ↑ of missingness and ↓ of branch lengths) |
|  | Under-merging | 0.92 – 0.99 | NA | ↓↓ proportion of shared loci and variable sites<br>↑↑ proportion of missing data | ↓ bootstrap support for very high bCTs (low effect)<br>↓↓ branch lengths |  |

1. *Optimization of the iCT by identifying an “under-merging threshold”.* As a first step, the dataset is assembled with a range of iCT (and a standard bCT). We employed a wide range of iCT values, but it can be narrowed as intra-specific diversity is expected to remain relatively low (e.g., iCTs≤0.85 are not expected to be biologically realistic in most cases). Our results suggest that the “under-merging threshold” might be identifiable simply by examining changes in cluster-related metrics. While this might be an interesting option when assembling large data sets, for which the between-samples clustering step could be time- and resource-consuming, we advocate that it is better to also compare the loci assemblies to ensure concordance between both groups of metrics. Once the threshold value has been identified, we suggest selecting the highest possible iCT value below it.
2. *Optimization of the bCT to maximize the informativeness of the assemblies while limiting the proportion of missing data.* Clusters built with the “optimal” iCT are subsequently used to produce assemblies with a range of bCTs. The range of values to be tested will depend on prior knowledge on the crown age of the focal taxa. We found that identifying an “under-merging threshold” was less straightforward here than for the iCT, but that a noticeable increase in the proportion of missing data, and a sharp contraction of branch lengths in concatenation trees were good indicators of under-merging. However, in case this threshold cannot be safely identified, it might be worth considering several alternative bCT values for downstream inferences.

These two steps are implemented in optiRADCT, a series of scripts enabling RADseq data set assembly with various iCTs and bCTs and visualisation of the results, available at https://github.com/rancilhac/optiRADCT. Once the iCT and bCT have been optimized, further loci filtering strategies can be explored, especially regarding the proportion of missing data in the assemblies, which have received much attention in recent studies (e.g., Cerca et al. 2021; Hühn et al. 2022; Crotti et al. 2019).

### 4.2 Varying the Clustering Threshold has a limited impact on phylogenetic and MSC inferences

Given the effects of the iCT and the bCT on the assemblies, one might predict that sub-optimal clustering thresholds would impact downstream phylogenetic analyses in various aspects. Reduced informativeness and increased missingness induced by under-merging are likely to affect tree topology, support, and reduce branch lengths. On the other hand, over-merging paralogs could either increase support for incorrect relationships or increase phylogenetic uncertainty by introducing noise in the data set (Leaché et al. 2015a). However, in the present case, concatenation-based phylogenetic analyses yielded very consistent topologies across iCT and bCT values. In all genera, inter-specific relationships and species monophyly were recovered with full support, except for a few nodes that received lower support for very low (≤0.60) or high (≥0.96) bCTs, although the topology itself remained constant. These relationships were also consistent with those recovered from the reference HE data sets, apart from the position of *R. parvipalmata* relative to *R. temporaria* (which could be due to influences of incomplete lineage sorting, as highlighted by concatenation- vs. species-tree differences in the RADseq data). On the other hand, intra-specific relationships were more variable, but no clear relationship with either the iCT or the bCT could be established. Thus, this variation likely reflects genuine phylogenetic uncertainty at the “grey zone” between intra- and inter-specific levels, where gene tree heterogeneity is expected to be higher due to gene flow and ancestral allele sharing among populations. In contrast to the topology, we found that the branch lengths of the trees were strongly affected by the bCT, and to a lower extent by the iCT, with both parameters having distinct effects. On the one hand, using greater iCTs induced a gradual, near-linear increase of the branch lengths, without any clear break useful for iCT optimization. On the other hand, increasing the bCT resulted in shorter branches, and the amplitude of the changes was much larger than that observed with the iCT. Branch length estimates are expected to vary according to several parameters, including the size and informativeness of the data set (Schwartz & Mueller 2010) and the proportion of missing data (Leaché et al. 2015b). Especially, in RADseq studies, using a higher Clustering Threshold will likely bias the assemblies toward slow-evolving loci, resulting in under-estimated branch lengths. Concordantly, we observe that branch length follows a near-linear relationship with the number of PIS and proportion of missing data for bCT>0.90. However, this trend does not hold for bCT<0.90, as the increase in branch lengths is not accompanied by an increase in the number of PIS, or a decrease in the proportion of missing data. In that case, overestimated branch lengths could result from the inclusion of paralogs due to loci over-merging.

Topological stability and node support in concatenation-based trees have often been used to select a supposedly correct CT for assembling RADseq data sets (e.g., Leaché et al. 2015a; McCartney-Melstadt et al. 2019). However, for our data sets, the recovered topology was resilient to changes in the iCT and bCT, with only unrealistically extreme values (e.g., ≤0.70 and ≥0.96) yielding lower support for some inter-specific relationships. It should be noted that this pattern might not be a general rule, as inferences in complex systems such as rapid radiations or genome-wide genealogical conflicts might be more sensitive to sub-optimal data assembly. Yet, our results suggest that topology and node support alone cannot always be used as accurate predictors of the quality of RADseq assemblies because, unlike branch lengths, they may often be largely unaffected by variation of assembly parameters.

As found in the concatenation-based analyses, the bCT did not have a dramatic effect on MSC inferences. In both *Discoglossus* and *Rana*, SNAPP recovered consistent and well supported topologies when using SNPs sets assembled with “optimal” or slightly “sub-optimal” bCTs (i.e., 0.88, 0.91 and 0.93). Using higher or lower values resulted in an increased uncertainty in the topology and parameter estimates, although the latter was difficult to assess considering the large confidence intervals. More strikingly, the analyses conducted with bCT=0.99 yielded noticeably shallower nodes, particularly close to the root. These results are consistent with the observed reduction of the concatenation trees’ branch lengths and might be an effect of the smaller size of the SNP matrices used, directly resulting from the loss of variability due to under-merging of loci. Furthermore, Bayes Factor species delimitation analyses consistently supported the same model ranking, regardless of the SNP matrix used. In both genera, “split” scenarios were slightly favoured compared to the current taxonomy, while “lump”scenarios always produced BF<1, a result consistent with recent studies showing that MSC species delimitation detects population structure rather than species limits (e.g., Sukumaran & Knowles 2017). However, the stable results recovered for both phylogenetic and MSC species delimitation analyses suggest that candidate species inferred from RADseq data based on either approach would be negligibly affected by suboptimal clustering threshold specifications.

## 5. Conclusions

Our results are encouraging for future RADseq-based phylogenomic studies. Indeed, other genome-wide data collection approaches, such as sequence capture, have often been preferred to RADseq for multiple reasons, including *de novo* assembly pitfalls (Leaché et al. 2015a; McKain et al. 2018; Hühn et al. 2022). We show here that RADseq and sequence capture data can yield equally accurate results, and that the clustering threshold is not a primary factor influencing the phylogenetic relationships inferred from the former. While unrealistic values, particularly those inducing strong under-merging, had an impact on the reconstructed relationships, moderate deviations from the “optimal” values produced well-supported topologies consistent with independent lines of evidence. Furthermore, in congruence with a previous study on plants (Clugston et al. 2019), we did not detect an obvious influence of genome size on RADseq data assembly and downstream phylogenetic inference, not even in newts with extremely large genomes. RADseq approaches have several advantages, including relatively low sequencing costs facilitating the generation of population-level data sets, and applicability to species without prior genomic knowledge. Hence, our data confirm RADseq as a promising as tool for accurate inference of evolutionary history and species limits in poorly known taxa, as observed in numerous recent studies (e.g., Razkin et al. 2016; Rodríguez et al. 2017; Paetzold et al. 2019; Rancilhac et al. 2019; Appelhans et al. 2020; Arrigoni et al. 2020; Dufresnes et al. 2020a, 2020b; Terraneo et al. 2021). However, it is of relevance that especially the bCT had a decisive impact on the estimation of branch lengths in concatenation trees, and that long and short branches were apparently affected differently. Thus, this parameter must be carefully considered for all downstream analyses relying on branch lengths, such as estimation of rates of character evolution, or divergence times (Leaché et al. 2015b; Collins & Hrbek 2018).

## Supporting information

Samples details

## Acknowledgments

This study was supported by the Deutsche Forschungsgemeinschaft (grant VE247/16-1 – HO 3492/6-1) in the framework of the ‘TaxonOmics’ priority program (SPP1991). We are gratefuk to M, Hofreiter (University of Potsdam) for valuable advice, and thank Glasgow Polyomics for assistance with RAD sequencing.

## Authors contributions

Conceptualization: LR, FS, CD, MV. Fieldwork and sampling: JWA, WB, PAC, GD, RD, PG, MPa, MPo, PP, JSP. HE data (Laboratory work and bioinformatics): CRH. RADseq data (Laboratory work and demultiplexing): MC, KE. Additional bioinformatics for the HE data, RADseq data assembly, phylogenetic analyses, analyses of RADseq assemblies, code writing: LR. Writing of the manuscript: LR, MV. Feedback and improvement of the manuscript: all authors. All authors have read the manuscript.

## Data availability statement

The supplementary material and appendix 1 are available at https://datadryad.org/stash/share/lyFID1l-7ZlgZNSNeFZWq_wYzYA0OnGFUrfCTgVzgjY. The raw RAD sequencing reads are available in the Short Read Archive under Bioproject <u>###### (to be added upon manuscript acceptance).</u> The alignments of the Hybrid-Enrichment markers, output of Ipyrad assemblies (assembly metrics, assembled loci and SNPs), as well as the concatenation and tetrad trees inferred from the RADseq data were uploaded to https://zenodo.org/record/7829243 (DOI:10.5281/zenodo.7829243). The code used to assemble the RADseq data, analyse the assemblies and produce plots is available at https://github.com/rancilhac/optiRADCT.

## References

Andrews, K. R., Good, J. M., Miller, M. R., Luikart, G., Hohenlohe, P. A. 2016. Harnessing the power of RADseq for ecological and evolutionary genomics. Nat. Rev. Genet., 17(2): 81–92.

Appelhans, M. S., Paetzold, C., Wood, K. R., Wagner, W. L. 2020. RADseq resolves the phylogeny of Hawaiian Myrsine (Primulaceae) and provides evidence for hybridization. J. Syst. Evol., 58(6): 823–840.

Arntzen, J. W., Wielstra, B., Wallis, G. P. 2014. The modality of nine *Triturus* newt hybrid zones assessed with nuclear, mitochondrial and morphological data. Biol. J. Linn., 113(2): 604–622.

Arrigoni, R., Berumen, M. L., Mariappan, K. G., Beck, P. S., Hulver, A. M., Montano, S., Pichon, M., Strona, G., Terraneo, T. I., Benzoni, F. 2020. Towards a rigorous species delimitation framework for scleractinian corals based on RAD sequencing: the case study of Leptastrea from the Indo-Pacific. Coral Reefs, 39(4), 1001–1025.

Bouckaert, R. R. 2010. DensiTree: making sense of sets of phylogenetic trees. Bioinformatics, 26(10): 1372–1373.

Bouckaert, R., Vaughan, T. G., Barido-Sottani, J., Duchêne, S., Fourment, M., Gavryushkina, A., Heled, J., Jones, G., Kühnert, D., De Maio, N., Matschiner, M., Mendes, F. K., Müller, N. F., Ogilive, H. A., du Pleissis, L., Popinga, A., Rambaut, A., Rasmussen, D., Siveroni, I., Suchard, M. A., Wu, C.-H., Xie, D., Zhang, C., Stadler, T., Drummond, A. J. 2019. BEAST 2.5: An advanced software platform for Bayesian evolutionary analysis. PLoS Comput. Biol., 15(4): e1006650.

Bryant, D., Bouckaert, R., Felsenstein, J., Rosenberg, N. A., RoyChoudhury, A. 2012. Inferring species trees directly from biallelic genetic markers: bypassing gene trees in a full coalescent analysis. Mol. Biol. Evol., 29(8): 1917–1932.

Capella-Gutiérrez, S., Silla-Martínez, J. M., Gabaldón, T. 2009. trimAl: a tool for automated alignment trimming in large-scale phylogenetic analyses. Bioinformatics, 25(15): 1972–1973.

Cariou, M., Duret, L., Charlat, S. 2013. Is RAD-seq suitable for phylogenetic inference? An in silico assessment and optimization. Ecol. Evol., 3(4): 846–852.

Cerca, J., Maurstad, M. F., Rochette, N. C., Rivera-Colón, A. G., Rayamajhi, N., Catchen, J. M., Struck, T. H. 2021. Removing the bad apples: A simple bioinformatic method to improve loci-recovery in de novo RADseq data for non-model organisms. Methods Ecol. Evol., 12(5): 805–817.

Chernomor, O., Von Haeseler, A., Minh, B. Q. 2016. Terrace aware data structure for phylogenomic inference from supermatrices. Syst. Biol., 65(6): 997–1008.

Clugston, J. A., Kenicer, G. J., Milne, R., Overcast, I., Wilson, T. C., Nagalingum, N. S. 2019. RADseq as a valuable tool for plants with large genomes—A case study in cycads. Mol. Ecol. Resour., 19(6): 1610–1622.

Collins, R. A., Hrbek, T. 2018. An in silico comparison of protocols for dated phylogenomics. Syst. Biol., 67(4), 633–650.

Cruaud, A., Gautier, M., Galan, M., Foucaud, J., Sauné, L., Genson, G., Dubois, E., Nidelet, S., Deuve, T., Rasplus, J.-Y. 2014. Empirical assessment of RAD sequencing for interspecific phylogeny. Mol. Biol. Evol., 31(5): 1272–1274.

Crotti, M., Barratt, C. D., Loader, S. P., Gower, D. J., Streicher, J. W. 2019. Causes and analytical impacts of missing data in RADseq phylogenetics: insights from an African frog (*Afrixalus*). Zool. Scr., 48(2), 157–167.

Degnan, J. H., Rosenberg, N. A. 2009. Gene tree discordance, phylogenetic inference and the multispecies coalescent. TREE, 24(6): 332–340.

Dufresnes, C., Nicieza, A. G., Litvinchuk, S. N., Rodrigues, N., Jeffries, D. L., Vences, M., Perrin, N., Martínez-Solano, Í. 2020a. Are glacial refugia hotspots of speciation and cytonuclear discordances? Answers from the genomic phylogeography of Spanish common frogs. Mol. Ecol., 29(5): 986–1000.

Dufresnes, C., Pribille, M., Alard, B., Gonçalves, H., Amat, F., Crochet, P.-A., Dubey, S., Perrin, N., Fumagalli, L., Vences, M., Martínez-Solano, I. 2020b. Integrating hybrid zone analyses in species delimitation: lessons from two anuran radiations of the Western Mediterranean. Heredity, 124(3): 423–438.

Eaton, D. A. 2014. PyRAD: assembly of de novo RADseq loci for phylogenetic analyses. Bioinformatics, 30(13): 1844–1849.

Eaton, D. A., Ree, R. H. 2013. Inferring phylogeny and introgression using RADseq data: an example from flowering plants (*Pedicularis*: Orobanchaceae). Syst. Biol., 62(5): 689–706.

Eaton, D. A., Overcast, I. 2020. ipyrad: Interactive assembly and analysis of RADseq datasets. Bioinformatics, 36(8): 2592–2594.

Edwards, R. J., Tuipulotu, D. E., Amos, T. G., O’Meally, D., Richardson, M. F., Russell, T. L., Vallinoto, M., Carneiro, M., Ferrand, N., Wilkins, M. R., Sequeira, F., Rollins, L. A., Holmes, E. C., Shine, R., White, P. A. 2018. Draft genome assembly of the invasive cane toad, *Rhinella marina*. Gigascience, 7(9): giy095.

Elshire, R. J., Glaubitz, J. C., Sun, Q., Poland, J. A., Kawamoto, K., Buckler, E. S., Mitchell, S. E. 2011. A robust, simple genotyping-by-sequencing (GBS) approach for high diversity species. PloS One, 6(5): e19379.

Fitch, W. M. 2000. Homology: a personal view on some of the problems. Trends Genet., 16(5): 227–231.

Gagnaire, P. A. 2020. Comparative genomics approach to evolutionary process connectivity. Evol. Appl., 13(6): 1320–1334

Goudarzi, F., Hemami, M. R., Rancilhac, L., Malekian, M., Fakheran, S., Elmer, K. R., Steinfartz, S. 2019. Geographic separation and genetic differentiation of populations are not coupled with niche differentiation in threatened Kaiser’s spotted newt (*Neurergus kaiseri*). Sci. Rep., 9(1): 6239.

Gregory, T.R. 2022. Animal Genome Size Database. [Accessed November 2020] http://www.genomesize.com

Hammond, S. A., Warren, R. L., Vandervalk, B. P., Kucuk, E., Khan, H., Gibb, E. A., Pandoh, P., Kirk, H., Zhao, Y., Jones, M., Mungall, A. J., Coope, R., Pleasance, S., Moore, R. A., Holt, R. A., Round, J. M., Ohora, S., Walle, B. V., Veldhoen, N., Helbing, C. C., Birol, I. 2017. The North American bullfrog draft genome provides insight into hormonal regulation of long noncoding RNA. Nat. Comm., 8(1), 1–8.

Herrera, S., Shank, T. M. 2016. RAD sequencing enables unprecedented phylogenetic resolution and objective species delimitation in recalcitrant divergent taxa. Mol. Phylogenet. Evol., 100: 70–79.

Hühn, P., Dillenberger, M. S., Gerschwitz-Eidt, M., Hörandl, E., Los, J. A., Messerschmid, T. F., Paetzold, C., Rieger, B., Kadereit, G. (2022). How challenging RADseq data turned out to favor coalescent-based species tree inference. A case study in *Aichryson* (Crassulaceae). Mol. Phylogenet. Evol., 167: 107342.

Hutter, C. R., Cobb, K. A., Portik, D. M., Travers, S. L., Wood Jr, P. L., Brown, R. M. 2022. FrogCap: A modular sequence capture probe-set for phylogenomics and population genetics for all frogs, assessed across multiple phylogenetic scales. Mol. Ecol. Resour., 22(3): 1100*–*1119.

Ilut, D. C., Nydam, M. L., Hare, M. P. 2014. Defining loci in restriction-based reduced representation genomic data from nonmodel species: sources of bias and diagnostics for optimal clustering. BioMed Res. Int., 2014: 675158.

Jombart, T., Dray, S. 2010. Adephylo: exploratory analyses for the phylogenetic comparative method. Bioinformatics, 26(15): 1–21.

Kalyaanamoorthy, S., Minh, B. Q., Wong, T. K., Von Haeseler, A., Jermiin, L. S. 2017. ModelFinder: fast model selection for accurate phylogenetic estimates. Nat. Methods, 14(6): 587–589.

Karbstein, K., Tomasello, S., Hodač, L., Dunkel, F. G., Daubert, M., Hörandl, E. 2020. Phylogenomics supported by geometric morphometrics reveals delimitation of sexual species within the polyploid apomictic *Ranunculus auricomus* complex (Ranunculaceae). Taxon, 69(6): 1191–1220.

Katoh, K., Standley, D. M. 2013. MAFFT multiple sequence alignment software version 7: improvements in performance and usability. Mol. Biol. Evol., 30(4): 772–780.

Keinath, M. C., Timoshevskaya, N., Timoshevskiy, V. A., Voss, S. R., Smith, J. J. 2018. Miniscule differences between sex chromosomes in the giant genome of a salamander. Sci. Rep., 8(1): 1–14.

Kuhner, M. K., Felsenstein, J. 1994. A simulation comparison of phylogeny algorithms under equal and unequal evolutionary rates. Mol. Biol. Evol., 11(3): 459–468.

Laurence, M., Hatzis, C., Brash, D. E. 2014. Common contaminants in next-generation sequencing that hinder discovery of low-abundance microbes. PloS One, 9(5): e97876.

Leaché, A. D., Fujita, M. K., Minin, V. N., Bouckaert, R. R. 2014. Species delimitation using genome-wide SNP data. Syst. Biol., 63(4): 534–542.

Leaché, A. D., Chavez, A. S., Jones, L. N., Grummer, J. A., Gottscho, A. D., Linkem, C. W. 2015a. Phylogenomics of phrynosomatid lizards: conflicting signals from sequence capture versus restriction site associated DNA sequencing. Genome Biol. Evol., 7(3): 706–719.

Leaché, A. D., Banbury, B. L., Felsenstein, J., De Oca, A. N. M., Stamatakis, A. 2015b. Short tree, long tree, right tree, wrong tree: new acquisition bias corrections for inferring SNP phylogenies. Syst. Biol., 64(6), 1032–1047.

Leaché, A. D., Bouckaert, R. R. 2018. Species trees and species delimitation with SNAPP: a tutorial and worked example. In Workshop on Population and Speciation Genomics, Český Krumlov.

Lemmon, A. R., Brown, J. M., Stanger-Hall, K., Lemmon, E. M. 2009. The effect of ambiguous data on phylogenetic estimates obtained by maximum likelihood and Bayesian inference. Syst. Biol., 58(1): 130–145.

Lemmon, E. M., Lemmon, A. R. 2013. High-throughput genomic data in systematics and phylogenetics. Ann. Rev. Ecol. Evol. Syst., 44: 99–121.

Mai, U., Mirarab, S. 2018. TreeShrink: fast and accurate detection of outlier long branches in collections of phylogenetic trees. BMC Genom., 19(5): 23–40.

Mastretta□Yanes, A., Arrigo, N., Alvarez, N., Jorgensen, T. H., Piñero, D., Emerson, B. C. 2015. Restriction site□associated DNA sequencing, genotyping error estimation and de novo assembly optimization for population genetic inference. Mol. Ecol. Resour., 15(1): 28–41.

McCartney-Melstad, E., Gidiş, M., Shaffer, H. B. 2019. An empirical pipeline for choosing the optimal clustering threshold in RADseq studies. Mol. Ecol. Resour., 19(5): 1195–1204.

McCormack, J. E., Hird, S. M., Zellmer, A. J., Carstens, B. C., Brumfield, R. T. 2013. Applications of next-generation sequencing to phylogeography and phylogenetics. Mol. Phylogenet. Evol., 66(2): 526–538.

McKain, M. R., Johnson, M. G., Uribe-Convers, S., Eaton, D., Yang, Y. 2018. Practical considerations for plant phylogenomics. Appl. Plant Sci., 6(3): e1038.

Miller, M. R., Dunham, J. P., Amores, A., Cresko, W. A., Johnson, E. A. 2007. Rapid and cost-effective polymorphism identification and genotyping using restriction site associated DNA (RAD) markers. Genome Res., 17(2), 240–248.

Mirarab, S., Warnow, T. 2015. ASTRAL-II: coalescent-based species tree estimation with many hundreds of taxa and thousands of genes. Bioinformatics, 31(12): i44–i52.

Nguyen, L. T., Schmidt, H. A., Von Haeseler, A., Minh, B. Q. 2015. IQ-TREE: a fast and effective stochastic algorithm for estimating maximum-likelihood phylogenies. Mol. Biol. Evol., 32(1): 268–274.

Paetzold, C., Wood, K. R., Eaton, D. A., Wagner, W. L., Appelhans, M. S. 2019. Phylogeny of Hawaiian *Melicope* (Rutaceae): RAD-seq resolves species relationships and reveals ancient introgression. Front. Plant Sci., 10: 1074.

Paradis, E., Claude, J., Strimmer, K. 2004. APE: analyses of phylogenetics and evolution in R language. Bioinformatics, 20(2): 289–290.

Paris, J. R., Stevens, J. R., Catchen, J. M. 2017. Lost in parameter space: a road map for stacks. Methods Ecol. Evol., 8(10): 1360–1373.

Peterson, B. K., Weber, J. N., Kay, E. H., Fisher, H. S., Hoekstra, H. E. 2012. Double digest RADseq: an inexpensive method for de novo SNP discovery and genotyping in model and non-model species. PloS One, 7(5): e37135.

Piwczyński, M., Trzeciak, P., Popa, M. O., Pabijan, M., Corral, J. M., Spalik, K., & Grzywacz, A. 2021. Using RAD seq for reconstructing phylogenies of highly diverged taxa: A test using the tribe Scandiceae (Apiaceae). J. Syst. Evol., 59(1): 58–72.

R Core Team 2020. R: A language and environment for statistical computing. R Foundation for Statistical Computing, Vienna, Austria. URL https://www.R-project.org/

Rambaut, A., Drummond, A. J., Xie, D., Baele, G., Suchard, M. A. 2018. Posterior summarization in Bayesian phylogenetics using Tracer 1.7. Syst. Biol., 67(5): 901–904.

Rancilhac, L., Goudarzi, F., Gehara, M., Hemami, M. R., Elmer, K. R., Vences, M., Steinfarz, S. 2019. Phylogeny and species delimitation of Near Eastern *Neurergus* newts (Salamandridae) based on genome-wide RADseq data analysis. Mol. Phylogenet. Evol., 133: 189–197.

Razkin, O., Sonet, G., Breugelmans, K., Madeira, M. J., Gómez-Moliner, B. J., Backeljau, T. 2016. Species limits, interspecific hybridization and phylogeny in the cryptic land snail complex *Pyramidula*: the power of RADseq data. Mol. Phylogenet. Evol., 101: 267–278.

Robinson, D. F., Foulds, L. R. 1981. Comparison of phylogenetic trees. Math. Biosci., 53(1-2), 131–147.

Rochette, N. C., Rivera-Colón, A. G., Catchen, J. M. 2019. Stacks 2: Analytical methods for paired-end sequencing improve RADseq-based population genomics. Mol. Ecol., 28(21): 4737–4754.

Rodríguez, A., Burgon, J. D., Lyra, M., Irisarri, I., Baurain, D., Blaustein, L., Göçmen, B., Künzel, S., Mable, B. K., Nolte, A. W., Veith, M., Steinfarz, S., Elmer, K. R., Philippe, H., Vences, M. 2017. Inferring the shallow phylogeny of true salamanders (*Salamandra*) by multiple phylogenomic approaches. Mol. Phylogenet. Evol., 115: 16–26.

Rogers, R. L., Zhou, L., Chu, C., Márquez, R., Corl, A., Linderoth, T., Freeborn, L., MacManes, M. D., Xiong, Z., Zheng, J., Guo, C., Xun, X., Kronsfort, M. R., Summers, K., Wu, Y., Yang, H., Richards-Zawacki, C. L., Zhang, G., Nielsen, R. 2018. Genomic takeover by transposable elements in the strawberry poison frog. Mol. Biol. Evol., 35(12): 2913–2927.

Rubin, B. E., Ree, R. H., Moreau, C. S. 2012. Inferring phylogenies from RAD sequence data. PloS One, 7(4): e33394.

Schmidt-Lebuhn, A. N., Aitken, N. C., Chuah, A. 2017. Species trees from consensus single nucleotide polymorphism (SNP) data: Testing phylogenetic approaches with simulated and empirical data. Mol. Phylogenet. Evol., 116: 192–201.

Schwartz, R. S., Mueller, R. L. 2010. Branch length estimation and divergence dating: estimates of error in Bayesian and maximum likelihood frameworks. BMC Evol. Biol., 10(1): 1–21.

Scornavacca, C., Delsuc, F., Galtier, N. 2020. Phylogenetics in the Genomic Era. Scornavacca, C.; Delsuc, F.; Galtier, N.. No commercial publisher | Authors open access book, p.p. 1–568, 2020, 978-2-9575069-0-3. ⟨hal-02535070v3⟩

Seppey, M., Manni, M., Zdobnov, E. M. 2019. BUSCO: assessing genome assembly and annotation completeness. In Gene prediction (pp. 227–245). Humana, New York, NY.

Smith, J. J., Putta, S., Zhu, W., Pao, G. M., Verma, I. M., Hunter, T., Bryant, S. V., Gardiner, D. M., Harkins, T. T., Voss, S. R. 2009. Genic regions of a large salamander genome contain long introns and novel genes. BMC Genom., 10: 1–11.

Stamatakis, A. 2014. RAxML version 8: a tool for phylogenetic analysis and post-analysis of large phylogenies. Bioinformatics, 30(9): 1312–1313.

Sukumaran, J., Knowles, L. L. 2017. Multispecies coalescent delimits structure, not species. Proc. Natl. Acad. Sci., 114(7): 1607–1612.

Sun, Y. B., Xiong, Z. J., Xiang, X. Y., Liu, S. P., Zhou, W. W., Tu, X. L., Zhong, L., Wang, L., Wu, D.-D., Zhang, B.-L., Zhu, C.-L., Yang, M.-M., Chen, H.-M., Li, F., Zhou, L., Feng, S.-H., Huang, C., Zhang, G.-J., Irwin, D., Hillis, D., Murphy, R. W., Yang, H.-M., Che, J., Wang, J., Zhang, Y.-P. 2015. Whole-genome sequence of the Tibetan frog *Nanorana parkeri* and the comparative evolution of tetrapod genomes. Proc. Natl. Acad. Sci., 112(11): E1257–E1262.

Terraneo, T. I., Benzoni, F., Arrigoni, R., Baird, A. H., Mariappan, K. G., Forsman, Z. H., Wooster, M. K., Bouwmeester, J., Marshell, A., Berumen, M. L. 2021. Phylogenomics of *Porites* from the Arabian Peninsula. Mol. Phylogenet. Evol., 161: 107173.

Zieliński, P., Nadachowska-Brzyska, K., Wielstra, B., Szkotak, R., Covaciu-Marcov, S. D., Cogălniceanu, D., Babik, W. 2013. No evidence for nuclear introgression despite complete mt DNA replacement in the Carpathian newt (*Lissotriton montandoni*). Mol. Ecol., 22(7): 1884–1903.

