## Supplementary materials for "Exploring the impact of read clustering thresholds on RADseq-based systematics: an empirical example from European amphibians"

**Supplementary S1: Detailed protocol for Hybrid-Enrichment data laboratory work and bioinformatic processing**

We used a modified version of the FrogCap sequence capture probe set (Hutter et al., 2021). In brief, our goal was to create a set of universal amphibian markers that could be targeted using a single set of probes. To initially select and design markers, we used the FrogCap markers (Hutter et al. 2021; GitHub: https://github.com/chutter/FrogCap-Sequence-Capture). We selected all FrogCap markers that were successfully captured broadly across Anura in the data from Hutter et al. (2021). Next, these markers were matched against the Salamander *Ambystoma* genome (Keinath et al. 2018) and markers were retained if they matched with at least 65% similarity. The total number of markers was 7,720 (UCEs: 2,122; exons: 5,598). We further aimed to add additional universal markers that could potentially be compared and included in phylogenetic analyses with other groups of organisms. We selected the USCO set of markers, which are orthologous genes found across deep evolutionary scales (i.e., Kingdom, Phylum, Class). To identify USCO markers present across Amphibia, we selected the salamander *Ambystoma mexicanum* and frogs *Xenopus tropicalis*, *Rana catesbiana*, *Nanorana parkeri*, *Rhinella marina*, and *Oophaga pumilio* genomes (Edwards et al. 2018; Hammond et al. 2017; Keinath et al. 2018; Rogers et al. 2018; Sun et al. 2015). We used the program BUSCO (v. 3.0.2; Seppey et al. 2019) to locate and identify the Metazoan and Tetrapoda USCO markers in these genomes. We included all Metazoan genes found in all the genomes, which totaled 677 out of 978 genes. We also added 29 genes from the Tetrapoda set to utilize remaining probes.

To create baits from the FrogCap markers, a consensus sequence was created from the alignments of the selected markers, as this has been suggested to improve target capture success across large phylogenetic scales (Hutter et al. 2021). For the USCO markers, we extracted each gene from each genome, and separated each of the genes into exons based on the boundaries determined by each genomes’ annotations. We aligned each exon and created a majority consensus sequence for that exon from across all the genomes. The consensus sequences were used to design a MYbaits-2 (40,040 baits) custom bait library (Arbor Biosciences), using 120mer baits to best capture sequences with greater than 5% divergence from the probes. The finalized set of markers were separated into probe sequences following a 2x tiling scheme, starting 20bp behind the start codon of the exon and tiling 120bp probes every 60bp until 20bp past stop codons. Individual probes were filtered using these criteria: (1) excluded probes that matched 70% length or greater to multiple locations in the genomes above with BLAST with a 70% similarity; (2) kept probes with a GC content between 30–50%; (3) kept if there were no repetitive sequences using RepeatMasker annotations from the online server (Sun et al. 2015); and (4) kept if there were no matches with BLAST to other probes (using a 70% similarity criterion). After filtration 40,211 baits remained; whole genes from the Tetrapoda set and their baits were randomly selected to total 189 baits from 7 genes to drop from the dataset to fit into the 40,040 bait limit. This resulted in a final set of 8,720 markers covering a total of 2,702,951 bp. The USCO markers totaled 14,153 baits (Metazoan: 13,196; Tetrapoda: 957) while the FrogCap markers totaled 25,884 baits (UCEs: 3,844; exons: 22,040).

Genomic DNA was extracted from the tissue samples using a PromegaTM Maxwell bead extraction robot and was quantified using a Promega QuantusTM fluorometer. Approximately 500 ng total DNA was acquired and set to a volume of 50 ul through dilution (with H20) or concentration (using a vacuum centrifuge) of the extraction when necessary. The genomic libraries for the samples were prepared by Arbor BioSciences library preparation service. Prior to library preparation, the genomic DNA samples were quantified using a Qubit and up to 4 µg was then taken to sonication with a QSonica Q800R instrument. After sonication and SPRI bead-based size-selection to modal lengths of roughly 300 bp, up to 500 ng of each sheared DNA sample were taken to Illumina Truseq-style sticky-end library preparation. Following adapter ligation and fill-in, each library was amplified for 6 cycles using unique combinations of i7 and i5 indexing primers, and then quantified with a Qubit. For each capture reaction, 125 ng of 8 libraries were pooled, and subsequently enriched for targets using the MYbaits v 3.1 protocol. Enrichment incubation times ranged 18–21 hours. Following enrichment, library pools were amplified for 10 cycles using universal primers and subsequently pooled in equimolar amounts for sequencing. Samples were sequenced on an Illumina HiSeq X with 150 bp paired-end reads in a shared lane with 96 total samples.

Illumina sequence data were de-multiplexed using the Illumina software bcl2fastq. Next, raw reads were cleaned of adapter contamination, low complexity sequences, and other sequencing artifacts using the program FASTP (default settings; Chen et al. 2018). Adapter-cleaned reads were then matched to a database of potential contaminants (human skin, ultra-pure water contamination, and other common bacteria; Laurence et al. 2014) and other genomes (*C. elegans*, *Drosophilia*) to ensure that no contamination persisted in our final dataset (see Hutter et al. 2021, Supplementary Table S3 for GenBank Accession Numbers of reference genomes). We decontaminated the adapter-cleaned reads with the program BBMAP from BBTools (https://jgi.doe.gov/data-and-tools/bbtools/), by matching cleaned reads to each reference contaminant genome (reads removed if they matched >95 percent similarity). The bioinformatics pipeline for filtering adapter contamination, assembling contigs, and exporting alignments is available at (https://github.com/chutter/FrogCap-Sequence- Capture).

Prior to assembly, the cleaned reads were further processed to decrease computational load and increase accuracy. We merged paired-end reads using BBMerge (Bushnell et al. 2017) from BBTools. BBMerge also fills in missing gaps between non-overlapping paired-end reads by assembling missing data from the other paired-end reads using the “Tadpole” program. Next, exact duplicates were removed if both read pairs were duplicated, using “dedupe” from BBTools. Additionally, duplicates from the set of merged paired-end contigs were removed if they were exact duplicates or were contained within another merged contig. Merged singletons and paired-end reads were assembled de novo using the program SPADES v.3.12 (Bankevich et al. 2012), which runs BAYESHAMMER (Nikolenko et al. 2013) error correction on the reads internally. Data were assembled using several different k-mer values (21, 33, 55, 77, 99, 127), in which orthologous contigs resulting from the different k-mer assemblies were merged. We used the dipSPAdes (Safonova et al. 2015) function to assemble contigs that were polymorphic by generating a consensus sequence from both haplotypes from orthologous regions such that polymorphic sites were resolved randomly.

Consensus haplotype contigs were then matched against reference marker sequences used to design the probes with BLAST (dc-megablast). Contigs were discarded if they failed to match ≥ 30% of the length of the reference marker, and contig matches fewer than 50 bp were removed. Contig matches to a given reference marker were discarded if more than one contig matched to the marker and were overlapping. For non-overlapping matches to the same reference marker, we merged these contigs by joining them together (Ns inserted in matching positions).

Next, our final set of matching markers was aligned on a marker-by-marker basis using MAFFT local pair alignment (settings: max iterations = 1000; ep = 0.123; op = 3; --adjust-direction; Katoh & Standley, 2013). We screened each alignment for samples ≥40% divergent from consensus sequences, which were almost always incorrectly assigned contigs. The obtained alignments were trimmed with trimAl (Capella-Gutiérrez et al., 2009), and subsequently re-aligned with MAFFT. Remaining poorly aligned regions were identified and trimmed using a custom R script. In brief, a consensus sequence was generated for each alignment, and the original sequences were divided in 80 bp long slices. The slices were then compared to the consensus sequence, and removed when >40% of the positions differed. Resulting alignments that were <40bp long or with less than 4 samples were filtered out. After trimming and filtering, the four data sets included 2,615 – 10,620 loci for a total of 921,033 – 6,751,735 bp (cf. Tab. S2 for more details).

**References**

Bankevich, A., Nurk, S., Antipov, D., Gurevich, A. A., Dvorkin, M., Kulikov, A. S., ... & Pevzner, P. A. (2012). SPAdes: a new genome assembly algorithm and its applications to single-cell sequencing. *Journal of computational biology*, *19*(5), 455-477.

Bushnell, B., Rood, J., & Singer, E. (2017). BBMerge–accurate paired shotgun read merging via overlap. *PloS one*, *12*(10), e0185056.

Capella-Gutiérrez, S., Silla-Martínez, J. M., & Gabaldón, T. (2009). trimAl: a tool for automated alignment trimming in large-scale phylogenetic analyses. *Bioinformatics*, *25*(15), 1972-1973.

Chen, S., Zhou, Y., Chen, Y., & Gu, J. (2018). fastp: an ultra-fast all-in-one FASTQ preprocessor. *Bioinformatics*, *34*(17), i884-i890.

Edwards, R. J., Tuipulotu, D. E., Amos, T. G., O'Meally, D., Richardson, M. F., Russell, T. L., ... & White, P. A. (2018). Draft genome assembly of the invasive cane toad, Rhinella marina. *Gigascience*, *7*(9), giy095.

Hammond, S. A., Warren, R. L., Vandervalk, B. P., Kucuk, E., Khan, H., Gibb, E. A., ... & Birol, I. (2017). The North American bullfrog draft genome provides insight into hormonal regulation of long noncoding RNA. *Nature Communications*, *8*(1), 1-8.

Hutter, C. R., Cobb, K. A., Portik, D. M., Travers, S. L., Wood Jr, P. L., & Brown, R. M. (2022). FrogCap: A modular sequence capture probe‐set for phylogenomics and population genetics for all frogs, assessed across multiple phylogenetic scales. *Molecular ecology resources, 22*(3): *1100–1119*.

Katoh, K., & Standley, D. M. (2013). MAFFT multiple sequence alignment software version 7: improvements in performance and usability. *Molecular biology and evolution*, *30*(4), 772-780.

Keinath, M. C., Timoshevskaya, N., Timoshevskiy, V. A., Voss, S. R., & Smith, J. J. (2018). Miniscule differences between sex chromosomes in the giant genome of a salamander. *Scientific reports*, *8*(1), 1-14.

Laurence, M., Hatzis, C., & Brash, D. E. (2014). Common contaminants in next-generation sequencing that hinder discovery of low-abundance microbes. *PloS one*, *9*(5), e97876.

Nikolenko, S. I., Korobeynikov, A. I., & Alekseyev, M. A. (2013, January). BayesHammer: Bayesian clustering for error correction in single-cell sequencing. In *BMC genomics* (Vol. 14, No. 1, pp. 1-11). BioMed Central.

Rogers, R. L., Zhou, L., Chu, C., Márquez, R., Corl, A., Linderoth, T., ... & Nielsen, R. (2018). Genomic takeover by transposable elements in the strawberry poison frog. *Molecular Biology and Evolution*, *35*(12), 2913-2927.

Safonova, Y., Bankevich, A., & Pevzner, P. A. (2015). dipSPAdes: assembler for highly polymorphic diploid genomes. *Journal of Computational Biology*, *22*(6), 528-545.

Seppey, M., Manni, M., & Zdobnov, E. M. (2019). BUSCO: assessing genome assembly and annotation completeness. In *Gene prediction* (pp. 227-245). Humana, New York, NY.

Sun, Y. B., Xiong, Z. J., Xiang, X. Y., Liu, S. P., Zhou, W. W., Tu, X. L., ... & Zhang, Y. P. (2015). Whole-genome sequence of the Tibetan frog Nanorana parkeri and the comparative evolution of tetrapod genomes. *Proceedings of the National Academy of Sciences*, *112*(11), E1257-E1262.

**Table S1.** Characteristics of the Hybrid-Enrichment data sets used as reference. Missingness is the percentage of the concatenation matrix represented by missing data. For loci length, the average, minimum and maximum values are given.

| Genus | Number of loci | Loci length (bp) | Concatenation length (bp) | Missingness (%) |
| --- | --- | --- | --- | --- |
| *Discoglossus* | 6,747 | 551.6 (80 – 5,082) | 3,708,712 | 31.73 |
| *Lissotriton* | 2,416 | 560.6 (80 – 2,056) | 1,354,496 | 19.66 |
| *Rana* | 10,620 | 628.5 (80 – 7,433) | 6,751,735 | 22.43 |
| *Triturus* | 2,615 | 359.6 (80 – 1,280) | 921,033 | 21.33 |

**Table S2.** Sizes of the SNPs matrices used for Multi-Species Coalescent analyses in SNAPP and BFD*.

| Genus | bCT=0.80 | bCT=0.85 | bCT=0.88 | bCT=0.91 | bCT=0.93 | bCT=0.95 | bCT=0.99 |
| --- | --- | --- | --- | --- | --- | --- | --- |
| *Discoglossus* | 265 | 278 | 275 | 272 | 261 | 234 | 54 |
| *Rana* | 652 | 673 | 691 | 687 | 660 | 632 | 108 |

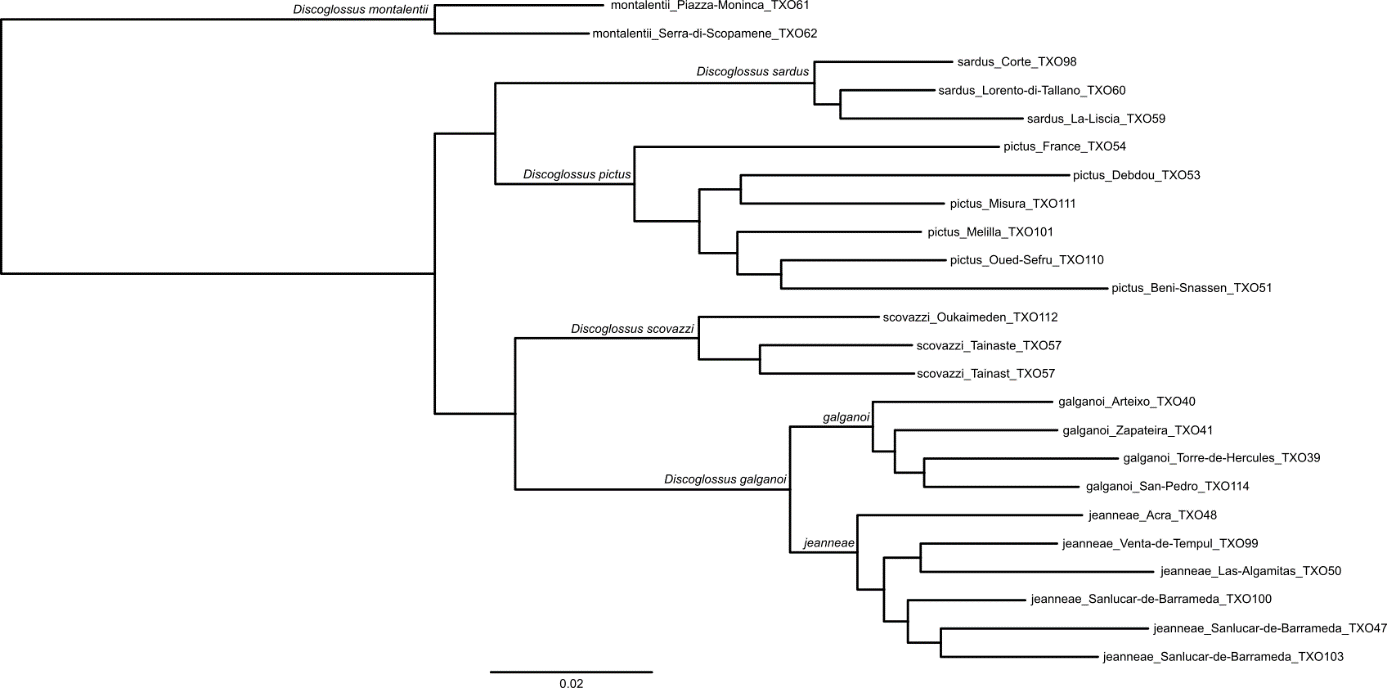

***Figure S1.*** *Complete Hybrid-Enrichment maximum-likelihood tree for the* Discoglossus *genus, inferred from a concatenation matrix of 3,708,712 bp. All nodes received an alRT support of 100.*

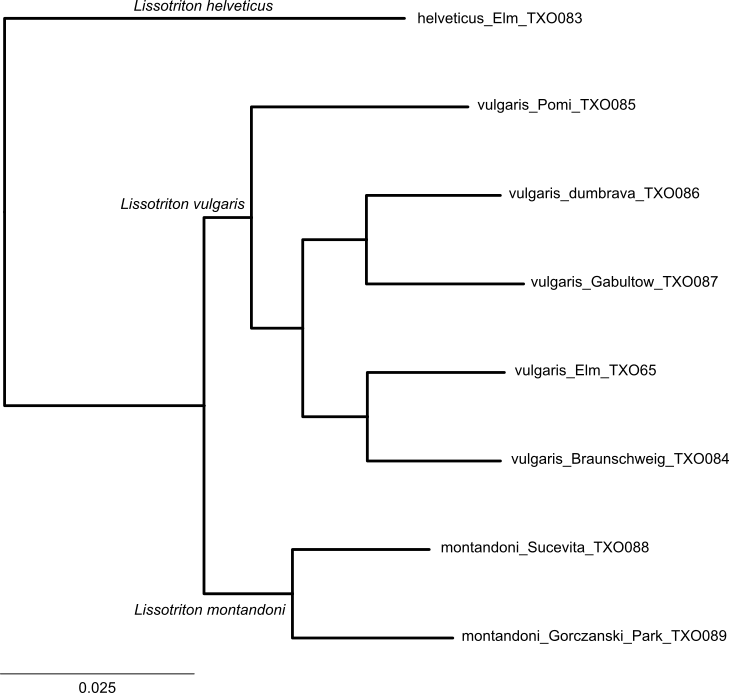

***Figure S2.*** *Complete Hybrid-Enrichment maximum-likelihood tree for the* Lissotriton *genus, inferred from a concatenation matrix of 1,354,496 bp. All nodes received an alRT support of 100.*

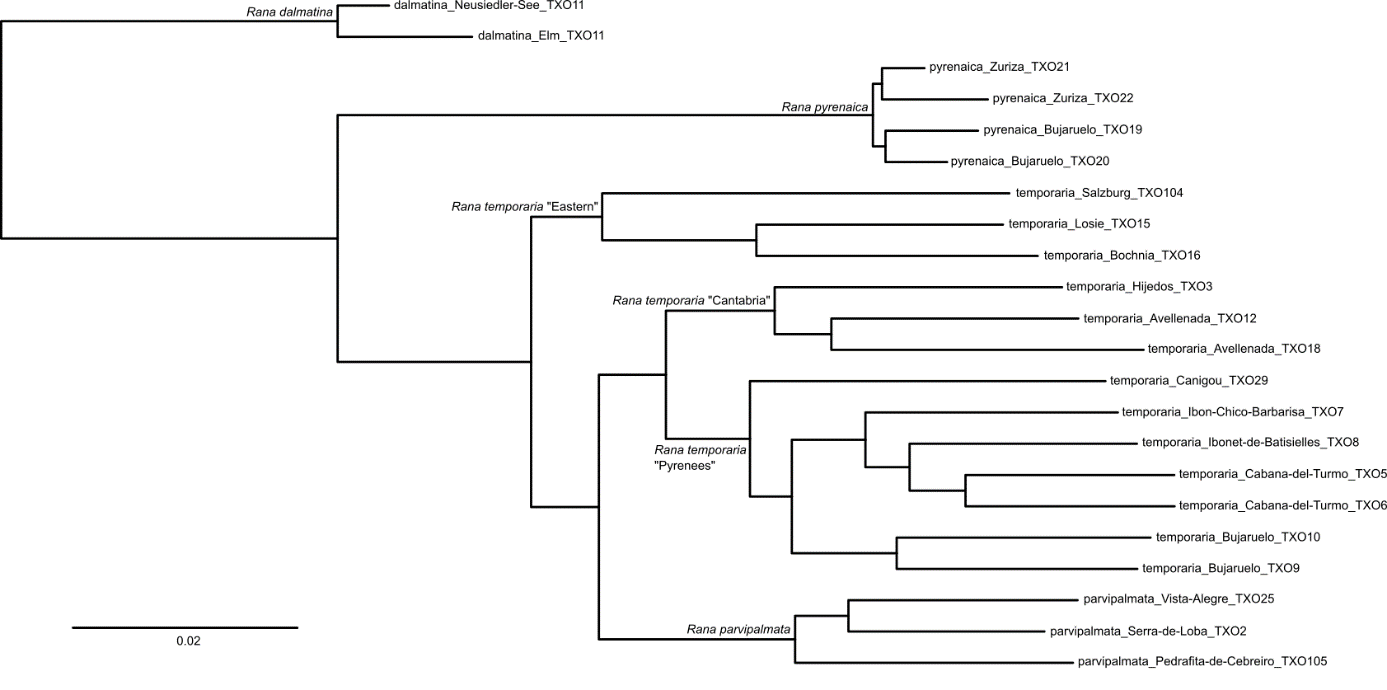

***Figure S3.*** *Complete Hybrid-Enrichment maximum-likelihood tree for the* Rana *genus, inferred from a concatenation matrix of 6,751,735 bp. All nodes received an alRT support of 100.*

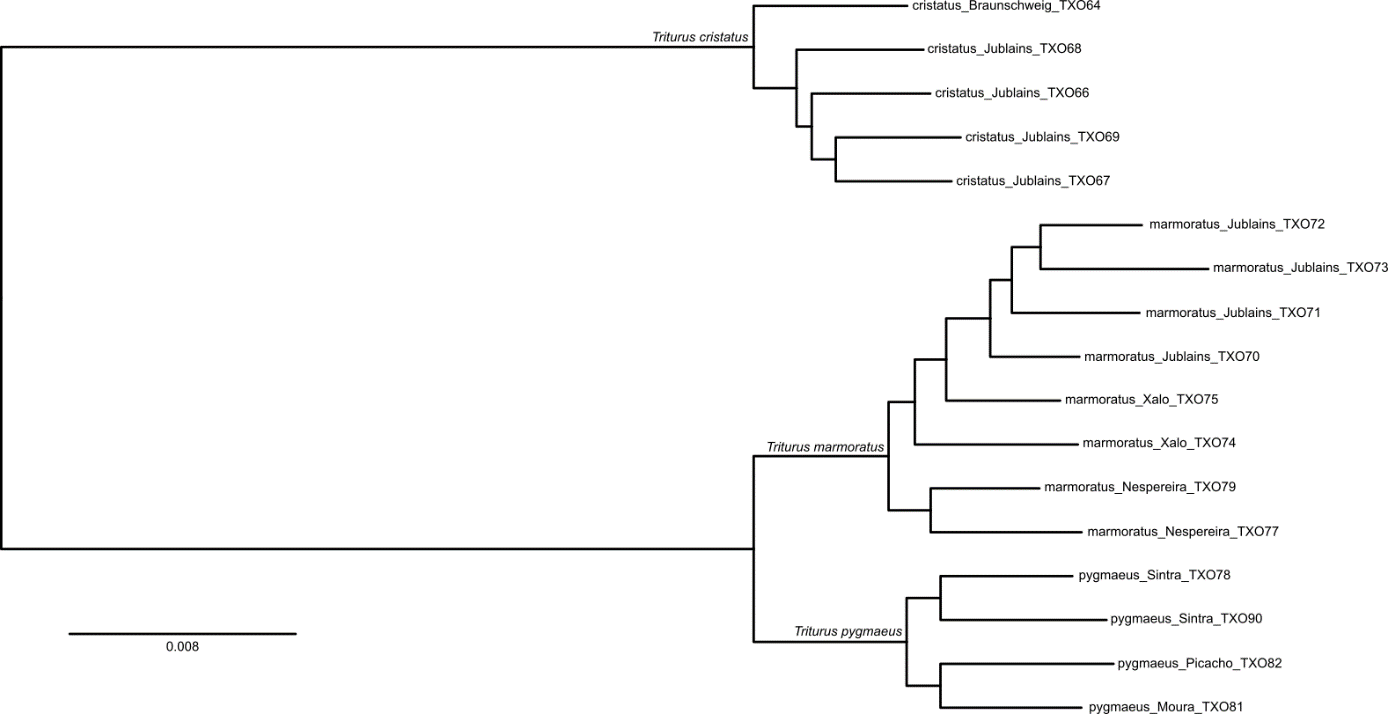

***Figure S4.*** *Complete Hybrid-Enrichment maximum-likelihood tree for the* Triturus *genus, inferred from a concatenation matrix of 921,033 bp. All nodes received an alRT support of 100.*

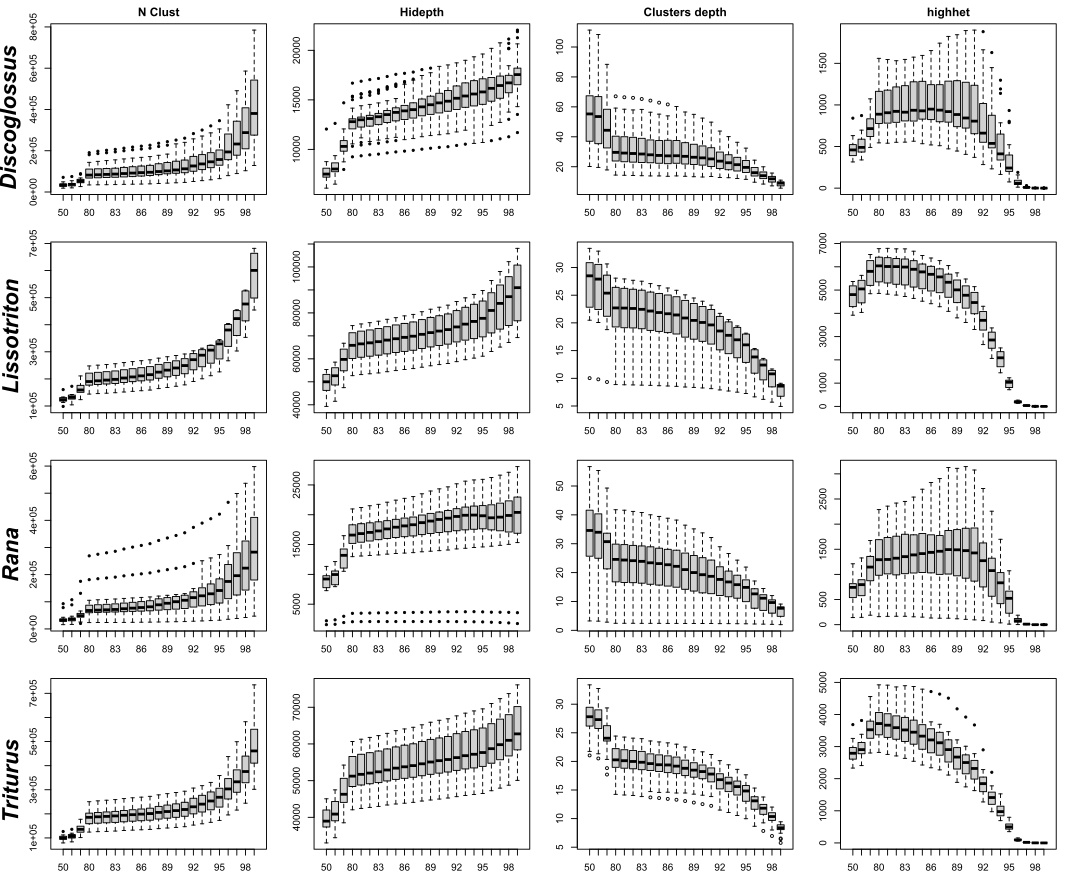

***Figure S5.*** *Change of clusters-related metrics as a function of the intra-samples clustering threshold (iCT). N Clust = Total number of clusters assembled; Hidepth = Number of clusters passing the depth filter; Clusters depth: number of reads in the clusters; Highhet = number of clusters rejected due to high heterozygosity. Note: the scale of the y-axis is not proportional to improve the visualisation of patterns at high iCTs.*

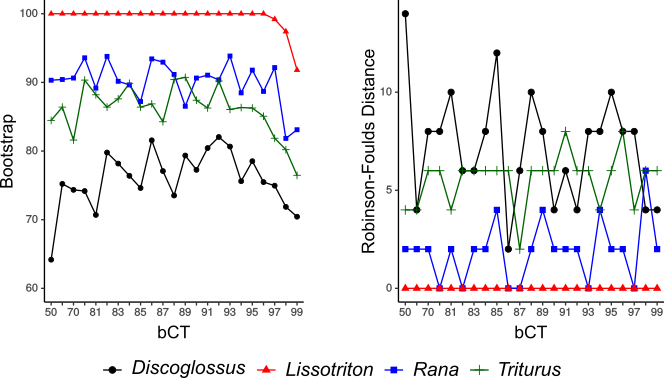

***Figure S6.*** *Effect of the between-samples Clustering Threshold (bCT) on Tetrad species tree inferences, represented as average bootstrap support (left) and topological distance to reference trees (right) as a function of the bCT.*

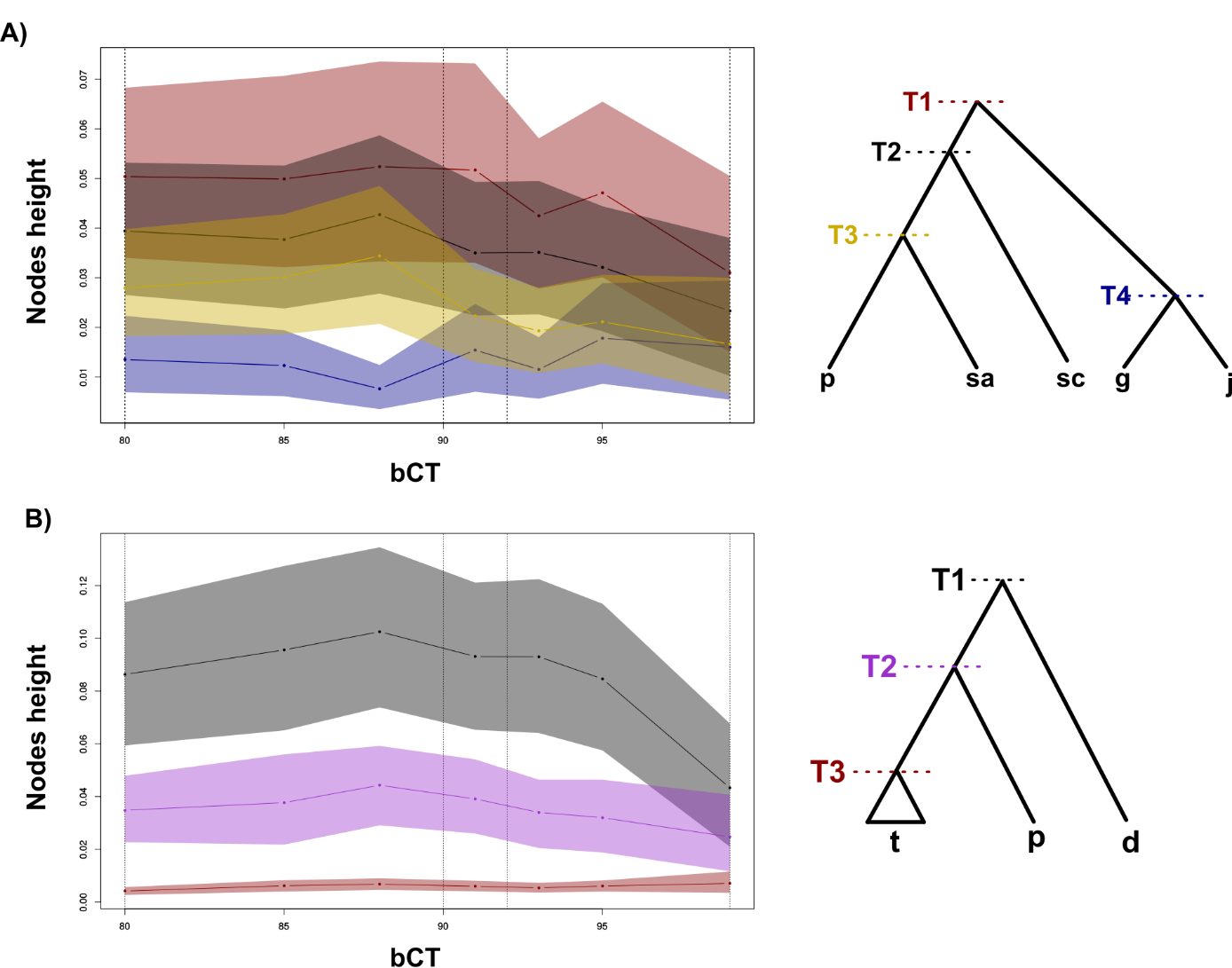

***Figure S7.*** *Effect of the between-samples Clustering Threshold (bCT) on node height estimates in SNAPP analyses in A)* Discoglossus *(g=*D. g. galganoi *; j=*D. g. jeanneae *; sc=*D. scovazzi *; sa=*D. sardus *; p=*D. pictus*) and B) Rana (d=*R. dalmatina *; p=*R. pyrenaica *; t=*R. temporaria *s.l., including* R. parvipalmata*). Color envelopes around the lines denote confidence intervals. Vertical dashed lines indicate the bCT value identified as optimal.*

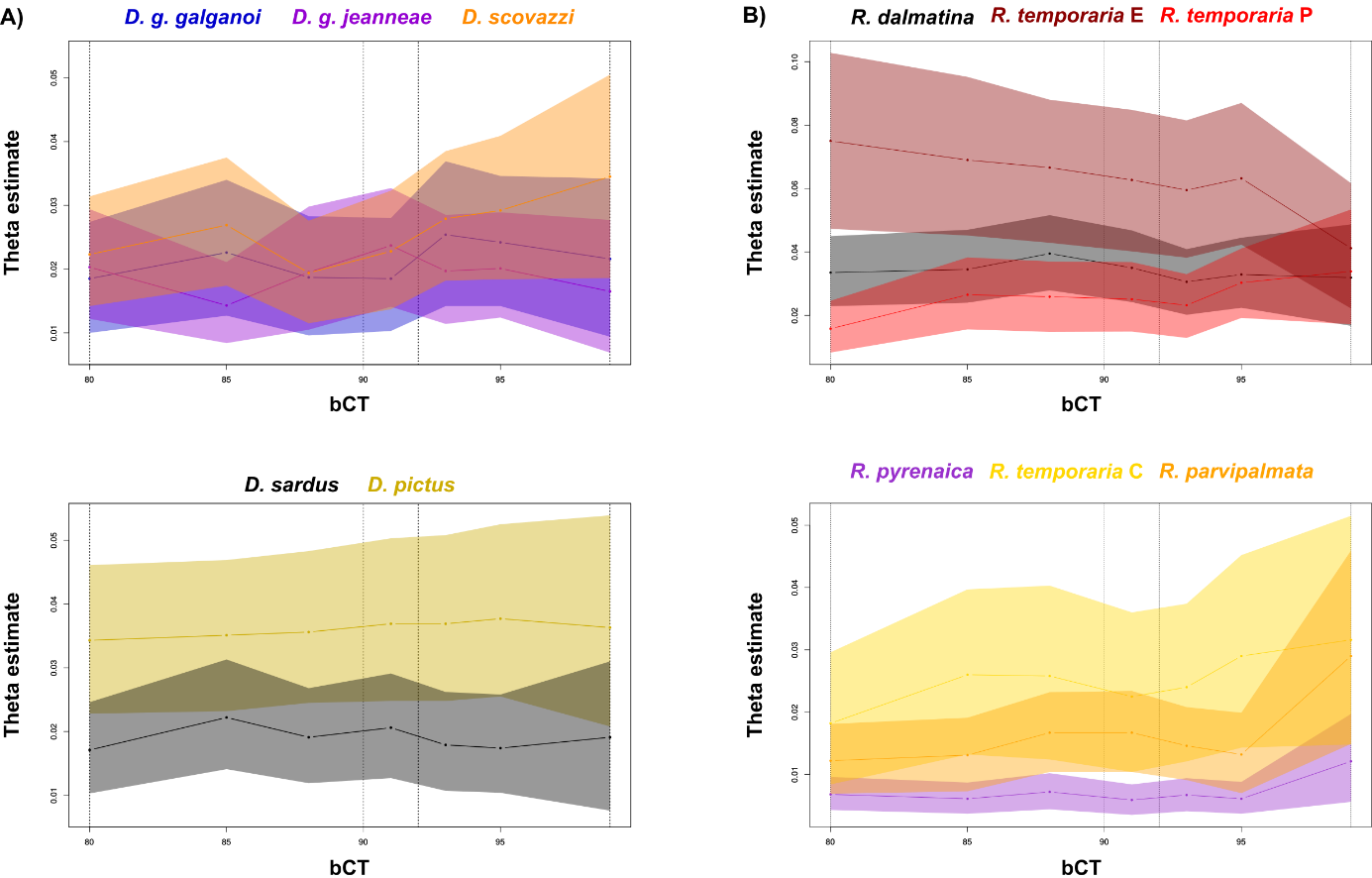

***Figure S8.*** *Effect of the between-samples Clustering Threshold (bCT) on population size (θ) estimates in SNAPP analyses in A) terminal branches in* Discoglossus *and B) terminal branches in* Rana*. Color envelopes around the lines denote confidence intervals. Vertical dashed lines indicate the bCT value identified as optimal.*
